# The Aging Microenvironment Shapes Alveolar Macrophage Identity in Aging

**DOI:** 10.1101/717033

**Authors:** Alexandra C. McQuattie-Pimentel, Ziyou Ren, Nikita Joshi, Satoshi Watanabe, Thomas Stoeger, Monica Chi, Ziyan Lu, Lango Sichizya, Raul Piseaux, Ching-I Chen, Saul Soberanes, Paul A. Reyfman, James M. Walter, Kishore R. Anekalla, Jennifer M. Davis, Kathryn A. Helmin, Constance E. Runyan, Hiam Abdala-Valencia, Kiwon Nam, Angelo Y. Meliton, Deborah R. Winter, Richard I. Morimoto, Gökhan M. Mutlu, Ankit Bharat, Harris Perlman, Cara J. Gottardi, Karen M. Ridge, Navdeep S. Chandel, Jacob I. Sznajder, William E. Balch, Benjamin D. Singer, Alexander V. Misharin, GR Scott Budinger

**Affiliations:** Department of Medicine, Division of Pulmonary and Critical Care Medicine, Northwestern University, Chicago, IL, 60611, USA; Department of Chemical and Biological Engineering, Northwestern University, Evanston, IL, 60208, USA; Department of Medicine, Division of Pulmonary and Critical Care Medicine, University of Chicago Hospitals, Chic; Department of Medicine, Division of Rheumatology, Northwestern University, Chicago, IL, 60611, USA; Department of Molecular Biosciences, Rice Institute for Biomedical Research, Northwestern University, Evanston, IL, 60208, USA; Department of Surgery, Division of Thoracic Surgery, Northwestern University, Chicago, IL, 60611, USA; The Scripps Research Institute Department of Chemical Physiology, La Jolla, CA, 10550, USA; Kanazawa University Graduate School of Medical Sciences, Department of Respiratory Medicine, Kanazawa, Ishikawa 920-8641, Japan

## Abstract

A dysfunctional response to inhaled pathogens and toxins drives a substantial portion of the susceptibility to acute and chronic lung disease in the elderly. We used transcriptomic profiling combined with genetic lineage tracing, heterochronic adoptive transfer, parabiosis and treatment with metformin to show that the lung microenvironment defines the phenotype of long-lived alveolar macrophages during aging. While tissue-resident alveolar macrophages persist in the lung without input from monocytes over the lifespan, severe lung injury results in their replacement with monocyte-derived alveolar macrophages. These monocyte-derived alveolar macrophages are also shaped by the microenvironment both during aging and in response to a subsequent environmental challenge to become transcriptionally and functionally similar to tissue-resident alveolar macrophages. These findings show that changes in alveolar macrophage phenotypes during injury and aging are not cell autonomous but instead are shaped by changes in the aging lung microenvironment.

## Introduction

Alveolar macrophages are derived from fetal monocytes that originate in the yolk sac or fetal liver. Shortly after birth these fetal monocytes rapidly differentiate into a stable, self-renewing population of “tissue-resident alveolar macrophages” (TRAM) that persist without input from bone marrow-derived monocytes for prolonged periods of time (Guilliams et al., 2013; Hashimoto et al., 2013; Janssen et al., 2011; Yona et al., 2013). TRAM reside in the ambient air and epithelial surface lining fluid, adhering to the lung epithelium via integrins on their surface where they are responsible for sampling, responding to, and clearing pathogens and particulates that reach the alveolar space (Watanabe et al., 2019). If TRAM are depleted then monocytes are recruited to the alveolar space where they differentiate into “monocyte-derived alveolar macrophages” (MoAM) through a progressive reshaping of their epigenome (Gibbings et al., 2015; Lavin et al., 2014).

Aging is the most important risk factor for the morbidity and mortality attributable to acute and chronic lung diseases (Budinger et al., 2017). Indeed, age-related lung diseases including pneumonia, Chronic Obstructive Pulmonary Disease and pulmonary fibrosis are among the most common causes of death in the developed world (WHO, 2018). Alveolar macrophages have been implicated as a key cell population in the pathobiology of all of these disorders (Hussell and Bell, 2014). The identification of ontologically distinct populations of TRAM and MoAM raises important questions about their function during aging. First, if TRAM persist in the alveolus without monocyte input over the lifespan, does their phenotype change with aging? Second, are age-related changes in TRAM cell-autonomous or driven by changes in the local microenvironment? Third, if MoAM are recruited to the lung in response to early life insults and persist, do they respond differently to a subsequent environmental challenge? To address these questions, we generated a genetic lineage tracing system to follow TRAM and MoAM over the lifespan and used transcriptomic identity to determine the role of the microenvironment and ontogeny in shaping their response to aging and repeated environmental stress. These results suggest that while ontogeny plays a major role in acute stress, its role in aging or chronic stress is minimal. Instead, our results highlight the importance of the lung microenvironment in shaping immune responses in the aging lung.

## Results

### Tissue-resident alveolar macrophages persist in the lung without input from bone marrow-derived monocytes during normal aging

To determine whether the ontogeny of TRAM changes during aging, we used thoracic shielding and a conditioning regimen that includes both radiation and systemic administration of busulfan to generate 4 month-old chimeric mice (CD45.1/CD45.2) with complete chimerism in monocytes and ∼80% preservation of TRAM (Misharin et al., 2017). We then allowed these animals to age without interventions up to 24 months (Figure 1A). If MoAM were less capable of self-renewal when compared with TRAM, the small population of MoAM recruited during the generation of chimeras would decline as a percentage of total cells during aging as MoAM are replaced by TRAM. However, both populations were stable over the lifespan (Figure 1B). Furthermore, MoAM showed comparable levels of EDU incorporation as TRAM (Figure 1C). These results suggest that MoAM and TRAM are similarly long-lived and capable of self-renewal, as has been previously reported (van de Laar et al., 2016). Nevertheless, the total number of alveolar macrophages was reduced in aged compared with young mice (Figure 1D).

**Figure 1.**
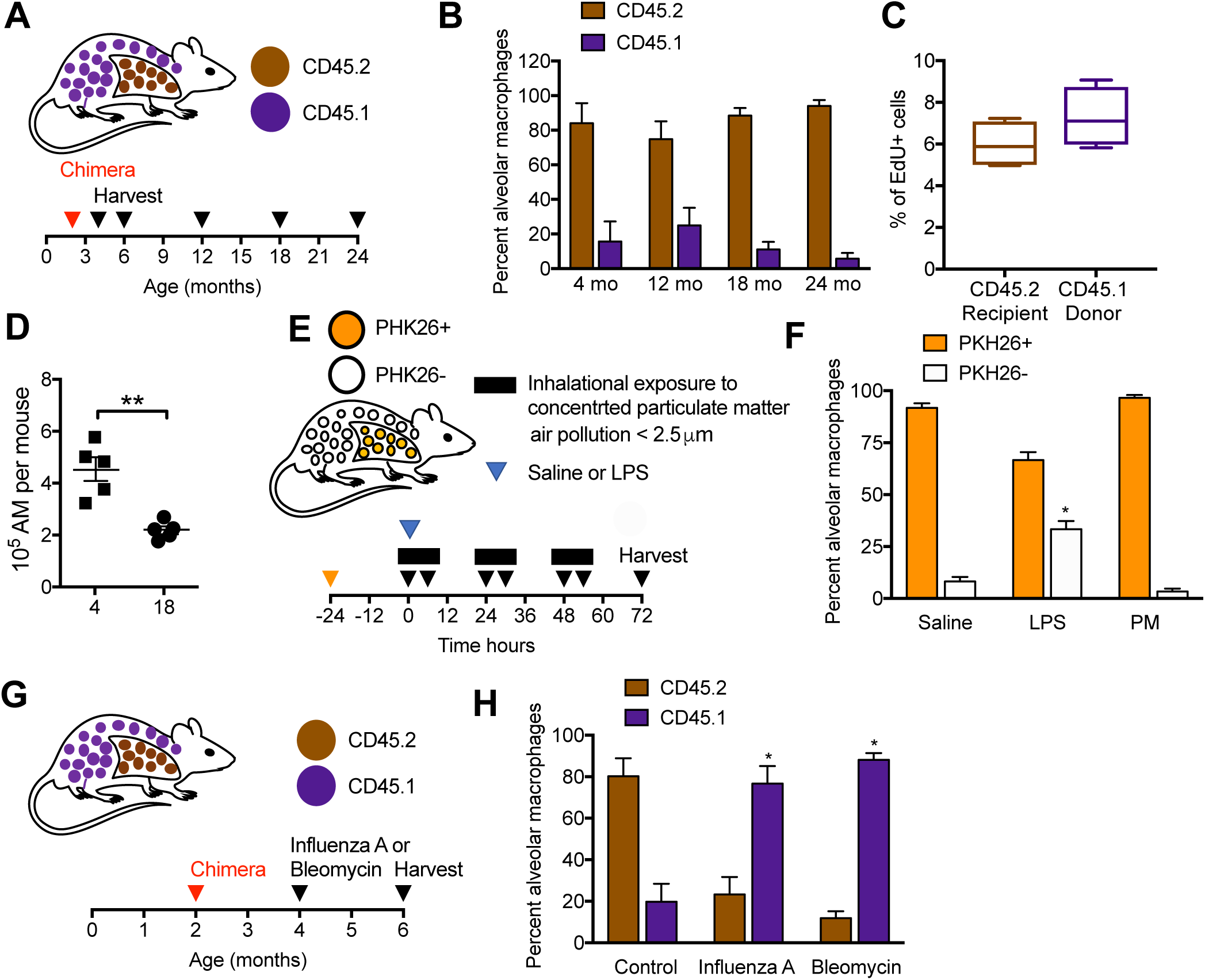
In the absence of severe injury, tissue-resident alveolar macrophages persist over the lifespan without input from bone marrow-derived monocytes. (A) Bone marrow chimeras with thoracic shielding were generated in which 100% of circulating monocytes were CD45.1 and ∼80% of tissue-resident alveolar macrophages were CD45.2. These mice were allowed to age in a barrier facility and harvested at the indicated times. (B) The percentage of monocyte-derived (CD45.1) or tissue-resident (CD45.2) alveolar macrophages over the lifespan was measured. Differences in the percentage of cells between ages were not significant (one-way ANOVA with Bonferroni correction, n=3-5 per time point). (C) Percentage of EdU+ cells in MoAM (CD45.1) and TRAM (CD45.2) one day after a single pulse of EDU was administered. Mice 4 months of age, n=4. (D) Total numbers of alveolar macrophages in young (4 months) and old (18 months) wild-type C57BL/6 mice. Student t-test, ** P = 0.001, n=5 mice per group. (E) Mice were intratracheally administered PKH26 fluorescent dye 24 hours before exposure to concentrated ambient particulate matter air pollution < 2.5 µm in diameter for 6 hours daily on three consecutive days. As negative and positive controls for recruitment of monocytes into the alveolar space mice received, PBS or LPS (1 mg/kg, intratracheally), respectively. See Figure S1 for gating strategy. (F) Percentage of alveolar macrophages that were labeled (PHK26+) or unlabeled (PHK26-) after PBS, LPS or exposure to concentrated ambient particulate matter air pollution (PM) via inhalation 6 hours daily for three consecutive weekdays. All mice were harvested after three days. One-way ANOVA with Bonferroni correction, n=5 per group, * indicates P<0.05 for comparison with control. (G) Shielded chimeric mice (4 months of age) were treated with influenza A virus or intratracheal bleomycin and then harvested 60 days later. (H) MoAM (CD45.1) and TRAM (CD45.2) were quantified by flow cytometry 60 days after infection with influenza A virus (A/WSN/33) or intratracheal administration of bleomycin. Control mice were not treated. * indicates P <0.05 for comparison with untreated mice.

Laboratory mice are housed in facilities where unusual steps are taken to limit exposure to inhaled pathogens and environmental particulates. In previous studies, investigators have shown that inhaled carbon-based particles are cleared by alveolar macrophages, which eventually are eliminated via mucociliary clearance (Semmler-Behnke et al., 2007). To determine whether this clearance of inhaled particulates is sufficient to induce the replacement of TRAM with MoAM, we exposed mice to concentrated urban particulate matter air pollution <2.5 µm in diameter at a dose we have previously reported to induce the release of IL-6 from alveolar macrophages and enhances the susceptibility to thrombosis (∼10 fold the concentration observed outside our laboratory in Chicago) for 6 hours per day on three consecutive days (Chiarella et al., 2014; Soberanes et al., 2019). We found that exposure to these typically encountered levels of airborne particles does not induce recruitment of MoAM to the lung, or deplete TRAM (Figure 1D and Fig. S1D). In contrast, the induction of severe lung injury with either intratracheal administration bleomycin or infection with sublethal dose of influenza A virus resulted in the recruitment of MoAM that persisted in the lung 60 days after the injury (Figure 1F,G). Notably, both bleomycin-induced lung injury and infection with influenza A virus induce rapid depletion of TRAM (Misharin et al., 2013).

### Aging is associated with reproducible changes in the transcriptome of tissue-resident alveolar macrophages driven by the microenvironment of the aging lung

Having established that TRAM are not replaced by MoAM over the lifespan, we explored whether TRAM change at the level of the transcriptome during normal aging using RNA-Seq. From the same mice, we flow sorted alveolar epithelial type II cells (Fig. S1C), with which alveolar macrophages are known to interact. We found significant differences in the transcriptome of both TRAM and alveolar type II cells from 18 month-old compared with 4 month-old animals (Figure 2A), which were similar in multiple independent cohorts of mice and were similar to transcriptomic changes in aging from another group (Wong et al., 2017) (Figure S2A-C). Consistent with previous studies of aging, immune system processes including genes related to interferon signaling were upregulated in aged compared with young animals, while genes related to cell cycle were downregulated (Figure 2A). To determine whether the transcriptomic changes in aged TRAM were cell-autonomous or were driven by factors in the local microenvironment, we performed heterochronic adoptive transfer of 1.5×10^5^ alveolar macrophages using 6 and 18 month-old mice as donors and recipients. Attempts to intratracheally adoptively transfer mature TRAM into untreated mice resulted in extremely poor engraftment (<5%) (Figure S2D-H), however, opening the alveolar niche by depleting TRAM via administration of liposomal clodronate before adoptive transfer allowed for engraftment rates up to 30% without the recruitment of neutrophils to the lung (Figure 2C, S2D-H). Interestingly, the rate of engraftment of young TRAM into older mice was significantly lower than the rate of engraftment of old TRAM into young mice (Fig. 2D,E). The overall structure of the transcriptomic data visualized by principal component analysis suggested that heterochronic transfer altered gene expression of adoptively transferred TRAM toward the age of the recipient (Fig. 2F,G). In contrast, heterochronic adoptive transfer of alveolar macrophages did not alter the transcriptome of alveolar type II cells (Fig. 2H,I). To exclude the possibility that the detected changes were due to the differences between CD45.1 and CD45.2 strains, we compared transcriptome of naïve TRAM from these strains and found that only 221 genes were differentially expressed (FDR q<0.05). Moreover, there was little overlap between genes differentially expressed with aging and those differentially expressed between TRAM from CD45.1 and CD45.2 mice (Fig. S2I, J).

**Figure 2.**
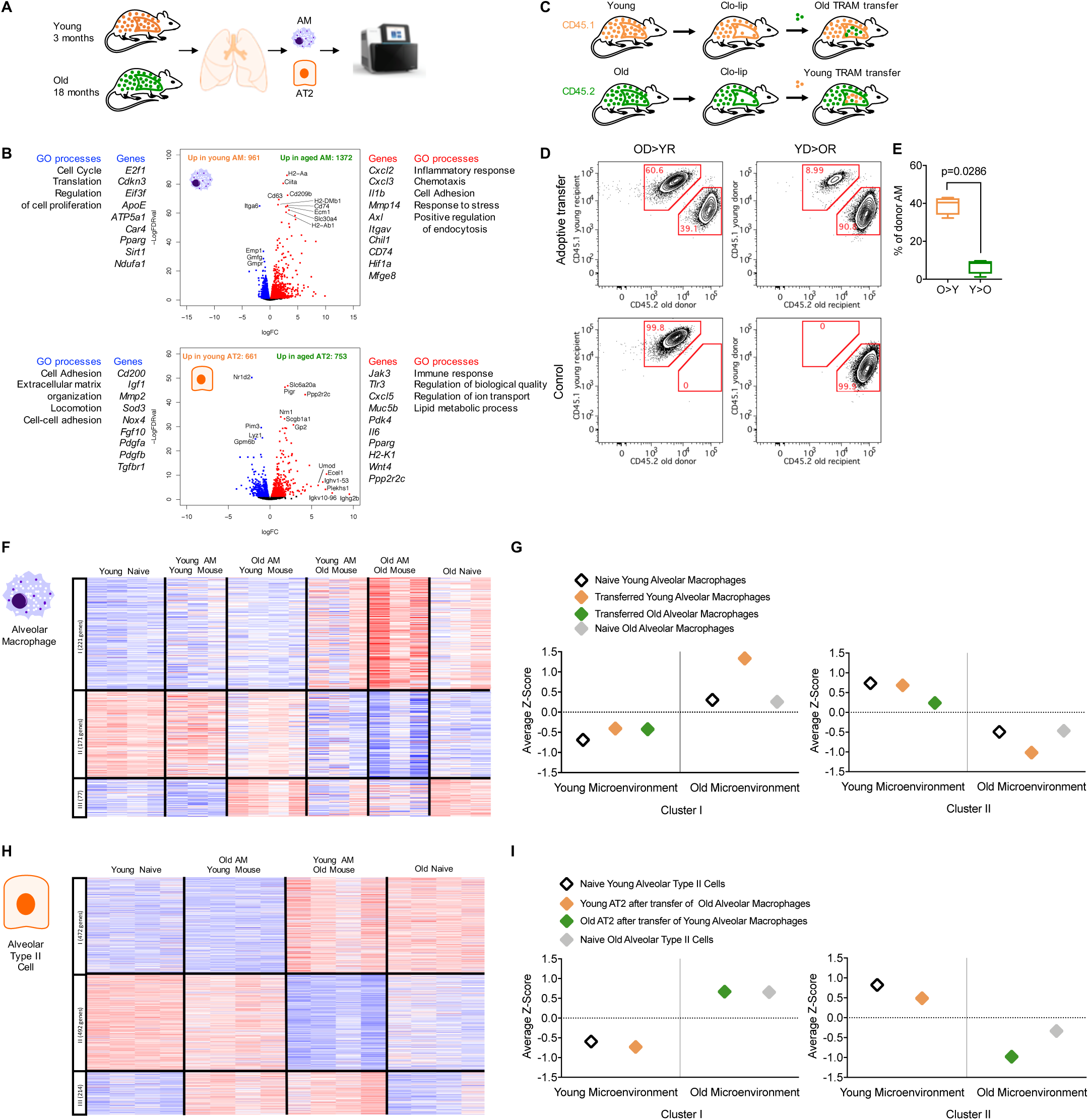
Age-related transcriptomic changes in tissue-resident alveolar macrophages are driven by the local microenvironment. (A) TRAM and alveolar epithelial type II cells were isolated via flow cytometry cell sorting from lung tissue from naïve C57Bl/6 mice (NIA NIH colony) at 4 (young) and 18 months (old) of age and analyzed using RNA-Seq (see also Figure S1C for genetic validation of the gating strategy for alveolar type II cell isolation). (B) Differentially expressed genes (FDR q<0.05) between TRAM and alveolar type II cells from young and old mice are shown with representative genes and GO biological processes (See Table S1 for complete list of genes and GO processes). (C) Heterochronic adoptive transfer experiments were performed using CD45.1/CD45.2 pairings as indicated (see also Figure S2). (D) Representative flow cytometry plots showing engraftment of TRAM from old and young donors into young and old mice, respectively. Harvest was performed 60 days after the adoptive transfer. All mice received liposomal clodronate (25 µL) intratracheally 72 hours prior to the adoptive transfer (also see Figure S2). (E) Percentage of engraftment for young and old TRAM, as in panel D, 4 mice per group. Mann-Whitney t-test. (F) Heatmap showing the results of k-means clustering of the differentially expressed genes (FDR q < 0.05 in ANOVA-like test) in TRAM 60 days after heterochronic adoptive transfer into young or old mice. Columns from left to right show (1) TRAM from young naïve mice, (2) TRAM in young recipient mice, (3) old TRAM adoptively transferred into young mice, (4) young TRAM adoptively transferred into old mice, (5) TRAM from old recipient mice, and (6) TRAM from naïve old mice (Figure S2 shows minimal overlap with differentially expressed genes in TRAM from CD45.1 compared with CD45.2 mice, see full list of genes in Table S2). (G) Average Z scores for the genes in Cluster I (genes up with aging) and Cluster II (genes down with aging) for each of the columns in (F). (H) Heatmap showing the results of k-means clustering of the differentially expressed genes (FDR q < 0.05 in ANOVA-list test) in alveolar type II cells 60 days after the heterochronic adoptive transfer of TRAM. Columns from left to right show transcripts in (1) alveolar type II cells from young naïve mice, (2) alveolar type II cells after adoptive transfer of young TRAM into old mice, (3) alveolar type II cells after adoptive transfer of old TRAM into old mice, and (4) alveolar type II cells from naïve old mice (see full list of genes in Table S2). Average Z scores for the genes in Cluster I (genes up with aging) and Cluster II (genes down with aging) for each of the columns in (H).

### Heterochronic parabiosis does not revert age-related changes in tissue-resident alveolar macrophages

Heterochronic parabiosis, in which circulating factors and myeloid cells from old or young mice are shared with age mismatched partners has long been recognized to reverse and induce age-related phenotypes in aged and young mice, respectively (Conboy et al., 2013). Accordingly, we used parabiosis to determine whether circulating factors or circulating cells from young animals (6 months of age) could reverse the environmentally-driven age-related transcriptomic changes in TRAM from old mice (18 months) (Figure 3A). We generated heterochronic and isochronic parabiont pairs and flow-sorted TRAM 60 days later, after parabionts established stable chimerism in the peripheral blood (Figure 3B). As alveolar type II cells express molecules necessary for the maintenance of TRAM, we simultaneously flow-sorted them and performed transcriptomic profiling via RNA-Seq (Figure 3H). Principal component analysis showed clustering of the samples according to age of the host irrespective of the age of the parabiont pair in both TRAM and alveolar type II cells (Figure 3C, 3I). Comparison of TRAM transcriptomes in young/young with old/old parabiont pairs revealed 866 differentially expressed genes (FDR < 0.05) (Figure 3E), 438 of which were also identified as differentially expressed in our independent aging dataset (Figure S3A, P=3.4 E-89). Comparison of TRAM in young/young pairings with TRAM from the aged environment in young/old pairings identified 474 differentially expressed genes, 288 of which overlapped with those differentially expressed during aging. In stark contrast, only 0 and 5 genes met criteria for differential expression between young/young and the young or old TRAM, respectively, from the heterochronic pairing (Figure 3F,G). Although the number of differentially expressed genes in alveolar type II cells was smaller (379), there was almost no change with heterochronic parabiosis (Figure 3I-M). These results suggest that transcriptional changes in TRAM and alveolar type II cells with aging are independent of circulating cell populations or signaling factors.

**Figure 3.**
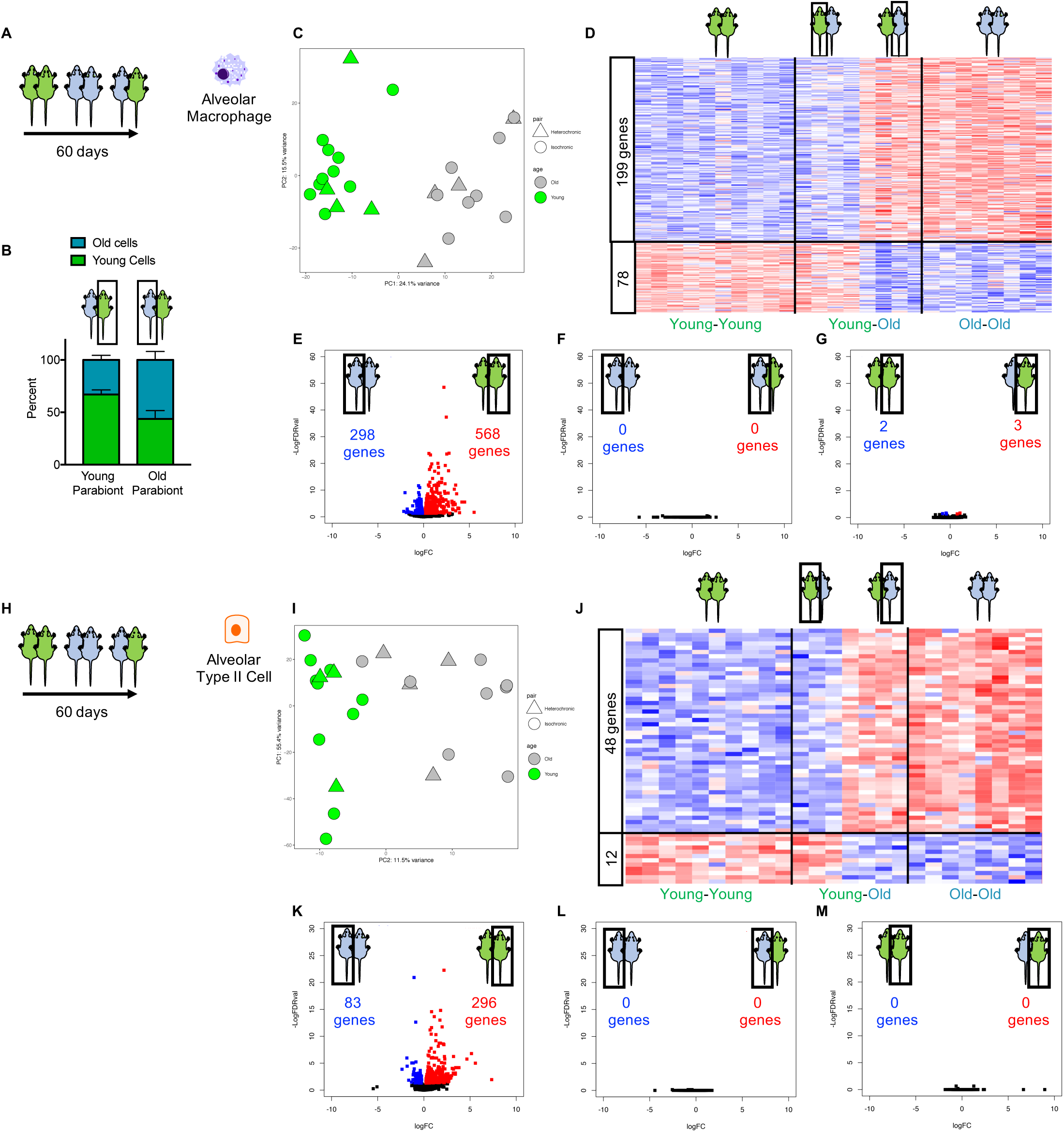
Heterochronic parabiosis does not reverse age-related transcriptomic changes in tissue-resident alveolar macrophages or alveolar type II cells. (A) Parabionts were generated from young (6 months, green) and aged (18 months, blue) pairs and TRAM were harvested after 60 days. (B) Percentage of circulating CD45^+^ cells from the young or old parabiont pair determined by flow cytometry using CD45.1/CD45.2. (C) PCA plot of TRAM harvested from young or aged mice linked to a isochronic or heterochronic parabiont pair. (D) Heatmap shows k-means clustering of differentially expressed genes inTRAM (FDR <0.01 in ANOVA-like test) between old and young mice with isochronic or heterochronic parabiont pairs (see also Table S4). (E) Volcano plot showing differentially expressed genes in TRAM from young/young versus old/old parabiotic pairs (FDR <0.05) (see also Figure S3 and Table S5). (F) Volcano plot showing differentially expressed genes in TRAM from old/old parabionts versus the old parabiont from young/old pairings (FDR <0.05) (see also Table S6). (G) Volcano plot showing differentially expressed genes in TRAM from young/young parabionts versus the young parabiont from young/old pairings (FDR <0.05) (see also Table S7). (H) Parabionts were generated from young and aged pairs and alveolar type II cells were harvested after 60 days. (I) PCA plot of alveolar type II cells harvested from young or old mice linked to a isochronic or heterochronic parabiont pair. (J) Heatmap shows k-means clustering of differentially expressed genes in alveolar type II cells (FDR <0.01 in ANOVA-like test) between old and young mice with isochronic or heterochronic parabiont pairs (see also Table S8). (K) Volcano plot showing differentially expressed genes in alveolar type II cells from young/young versus old/old parabiotic pairs (FDR <0.05) (see also Figure S3 and Table S9). (L) Volcano plot showing differentially expressed genes in alveolar type II cells from old/old parabionts versus the old parabiont from young/old pairings (FDR <0.05) (see also Table S10). (M) Volcano plot showing differentially expressed genes in alveolar type II cells from young/young parabionts versus the young parabiont from young/old pairings (FDR <0.05) (see also Table S11).

### Chronic administration of metformin induces only small changes in the aging transcriptome of tissue-resident alveolar macrophages and alveolar type II cells

Metformin has been shown to extend lifespan and improve healthspan in mice (Martin-Montalvo et al., 2013). We have previously reported that the short-term administration of metformin resulted in few changes in baseline gene expression in TRAM but enhanced the expression of chaperone after exposure to particulate matter air pollution (Soberanes et al., 2019). To determine whether chronic treatment with metformin could reverse age-related changes in gene expression in TRAM or alveolar type II cells, we treated young (4 month) or old (18 month) mice with metformin for 60 days in the drinking water and flow-sorted TRAM and alveolar type II cells for RNA-Seq (Figure 4B-G). We found that the administration of metformin induced relatively few changes in gene expression in TRAM from young or old mice (130 and 134 genes, respectively, FDR q < 0.05, Figure 4C-E). Despite the small number of genes, there was significant overlap with genes associated with aging (37 genes, P < 0.01 by hypergeometric test, Table S12) and these genes directionally moved toward a younger age (Table S12). The genes included some with potential biologic relevance to aging including *Il6ra*, *Cflar*, important for monocyte to macrophage differentiation, the calcium uniporter *Mcub*, and the cyclins *Cdk13* and *Ccng2*. In alveolar type II cells there were similar small changes in gene expression with metformin (116 and 79, respectively, Figure 4H-J). Of these, 37 of the 97 genes overlapped with those that changed during aging (including *Socs3*, *Junb*, and *Hmox1* (P=0.01)). These results suggest that while metformin has modest effects on basal gene expression it does alter the expression of some age-related genes.

**Figure 4.**
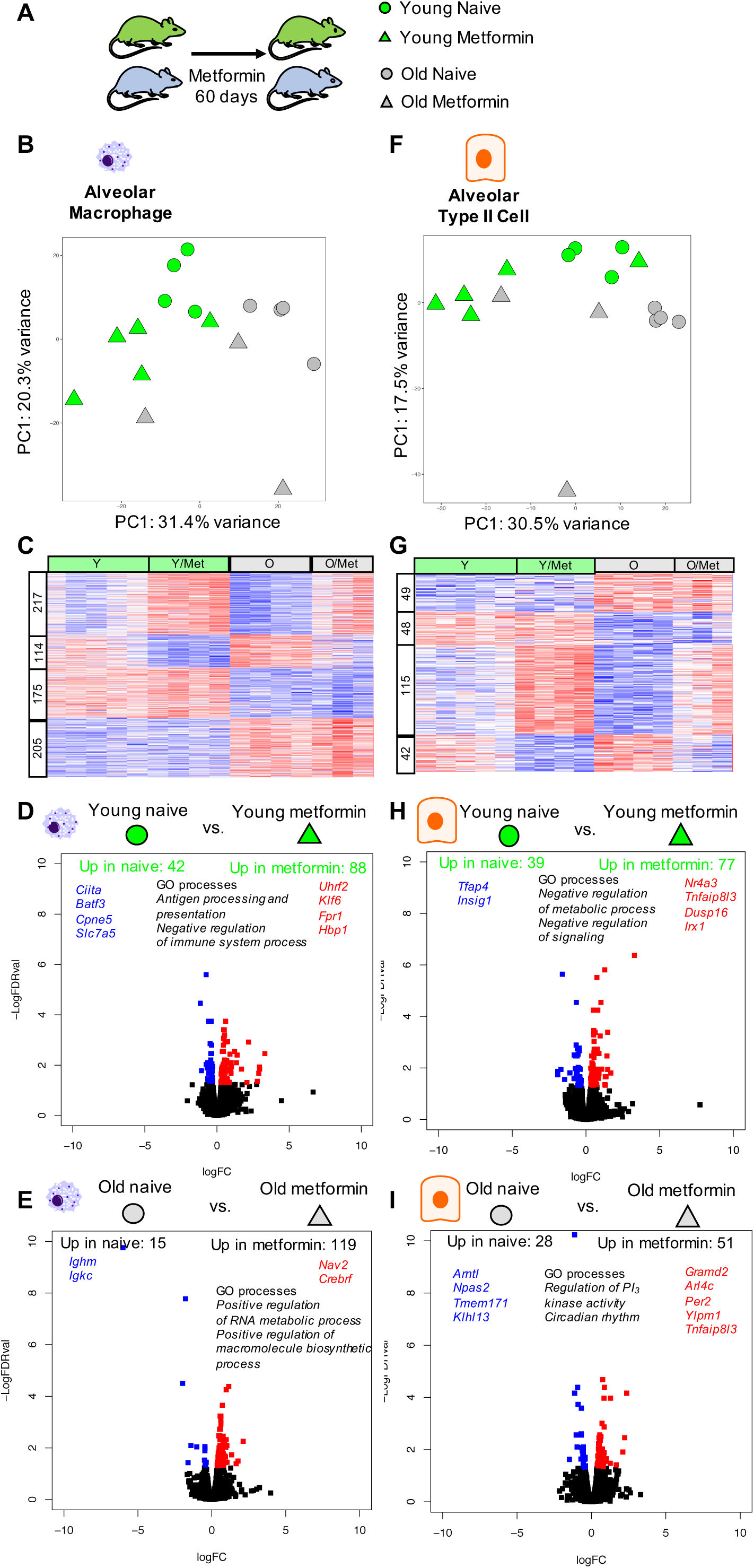
Metformin results in modest differences in age-related changes in the transcriptome of tissue-resident alveolar macrophages and alveolar type II cells. (A) Young (4 month) and old (18 month) mice were treated with metformin in the drinking water for 60 days and TRAM and alveolar type II cells were harvested for RNA-Seq. (B) PCA plot of transcriptomes of TRAM from animals in the four conditions. (C) Heatmap shows differentially expressed genes in TRAM from metformin treated animals (FDR <0.01 in ANOVA like test) (see also table S13). (D) Volcano plot shows differences in alveolar macrophages from young mice treated with metformin (FDR <0.05). Individual genes are highlighted, and GO processes are shown in the center (See table S14 for a full list of genes and enriched GO processes). (E) Volcano plot shows differences in TRAM from old mice treated with metformin (FDR <0.05). Individual genes are highlighted. (See table S15 for a full list of genes and enriched GO processes). (F) PCA plot of transcriptomes of alveolar type II cells from animals in the four conditions. (G) Heatmap shows differentially expressed genes in alveolar type II cells from metformin treated animals (FDR<0.01 in ANOVA like test) (see also Table S16). (H) Volcano plot shows differences in alveolar type II cells from young mice treated with metformin (FDR <0.05). Individual genes are highlighted, and GO processes are shown in the center. See Table S17 for a full list of genes and enriched GO processes. (I) Volcano plot shows differences in alveolar type II cells from old mice treated with metformin (FDR<0.05). Individual genes are highlighted (see Table S18 for a full list of genes and enriched biological GO processes).

### Differences between tissue-resident and monocyte-derived alveolar macrophages persist during aging

We took advantage of the small but stable chimerism in our shielded bone marrow chimeric mice to compare transcriptomic changes with aging in MoAM and TRAM, which were similar (Figure 5A). Nevertheless, a number of genes were differentially expressed over time, and, indeed, further diverged with aging (Cluster IV and V, Figure 5A). These MoAM were evenly distributed in the lung tissue (Figure 5B), thus, the observed differences in the transcriptome of MoAM and TRAM likely reflect their ontogeny. We observed significant overlap between these differentially expressed gene and genes we previously reported to be differentially expressed between TRAM and MoAM 10 months after bleomycin-induced lung injury (Misharin et al., 2017) (Figure 5C). Similarly we observed all of the genes identified as differentially expressed between MoAM and TRAM after liposomal clodronate using microarray technology (Gibbings et al., 2015). There was little overlap between these genes and those differentially expressed genes TRAM harvested from CD45.1 and CD45.2 mice (Figure S5). These results reveal consistent transcriptional differences between TRAM and MoAM residing in the same microenvironment irrespective of the stimulus that led to their recruitment. While several groups have associated histone modifications in the differentiation of monocytes to macrophages and in the development of innate immune memory, the importance of DNA methylation in this process has not been explored (Gosselin et al., 2014; Lavin et al., 2014; Morales-Nebreda et al., 2019; Netea et al., 2016; Quintin et al., 2012; Saeed et al., 2014; Yoshida et al., 2015). Accordingly, we performed reduced representation bisulfite sequencing on TRAM and MoAM from aged chimeric animals. We looked for evidence of differential DNA methylation in promoter regions 1000 bp upstream and downstream of the start site of differentially expressed genes in our RNA-Seq dataset and in putative enhancer regions specific to alveolar macrophages defined as H3K4me1 peaks identified by Lavin et al. exclusive of promoter regions (Figure 5D) (Lavin et al., 2014; Weinberg et al., 2019). We found no evidence of differences in these regions.

**Figure 5.**
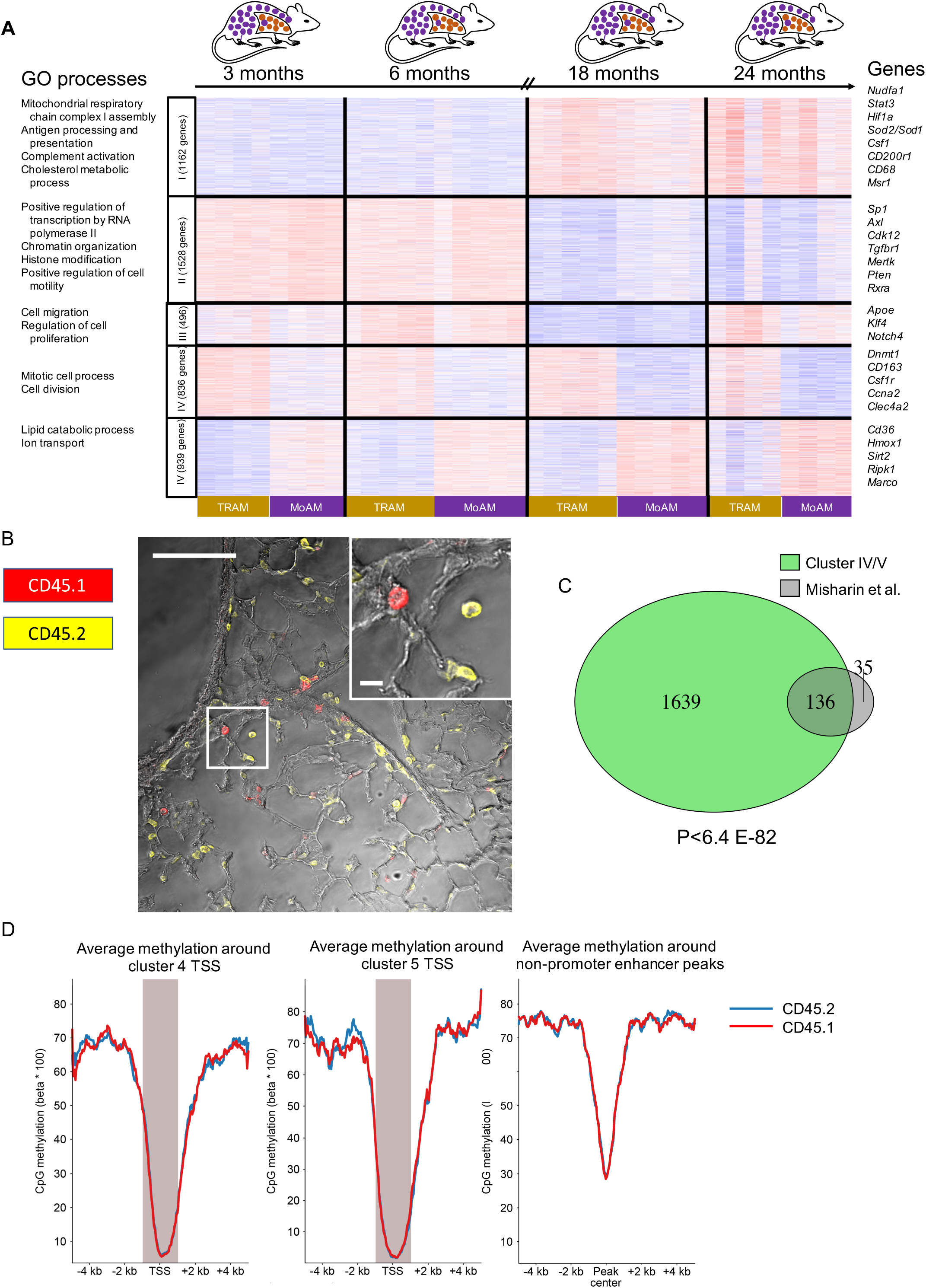
Transcriptional differences between monocyte-derived alveolar macrophages and tissue-resident alveolar macrophages persist over the lifespan. (A) Alveolar macrophages from shielded chimeric mice were harvested at the indicated ages and TRAM and MoAM were flow sorted based on the CD45.1 or CD45.2 label. Differentially expressed genes (FDR <0.01 in ANOVA-like test) were identified and subjected to k-means clustering. Selected genes and GO processes from each cluster are highlighted (see also Table S19 for full list of genes and GO processes). (B) Representative immunofluorescence image of a lung section from a shielded chimeric mouse (6 months of age) stained for CD45.2 to mark TRAM and CD45.1 to mark MoAM. Combined image overlaid on phase contrast image is shown. Scale bar is 10 µm. (C) Venn diagram showing overlap of differentially expressed genes between MoAM and TRAM in this model with an independent dataset (Misharin et al., 2017) collected 10 months after bleomycin exposure. (D) Reduced representation bisulfite sequencing was performed on TRAM and MoAM from 6 month-old mice. The frequency of methylated CpG motifs in promoter regions within 1000 bp upstream and downstream of the transcriptional start site of differentially expressed genes between TRAM and MoAM in shielded chimeric mice was compared with their frequency across the genome. A similar analysis was performed using putative enhancer regions specific to alveolar macrophages defined as consensus H3K4me1 peaks by (Lavin et al., 2014). No significant differences in DNA methylation were detected.

### Sequential injury models provide a physiologically relevant model to determine whether alveolar macrophage ontogeny affects the response to repeated environmental challenge in aging

The findings above suggest that in some individuals, episodic lung injury over the lifespan could result in at least two transcriptionally distinct populations of alveolar macrophages within the same microenvironment. We and others have reported that MoAM play a causal role in the development of bleomycin-induced lung fibrosis and influenza A virus-induced pneumonia (Kim et al., 2008; McCubbrey et al., 2018; Misharin et al., 2017). We therefore reasoned that the induction of ontological chimerism after infection with the influenza A virus or treatment with bleomycin (Figure 1G) would provide a physiologically relevant system to determine whether differential responses of MoAM and TRAM to a second stimulus underlie the observation of innate immunologic memory. Accordingly, we generated a cohort of shielded chimeric mice and subjected them to sequential injury models (Figure 6A,E). We harvested animals and performed flow cytometry sorting of TRAM and MoAM and analyzed them using RNA-Seq (Figure S6). Principal component analysis of the entire dataset demonstrates good reproducibility in the data and shows markedly different responses of MoAM and TRAM to influenza A virus-induced pneumonia and bleomycin-induced pulmonary fibrosis (Figure 6B,F).

**Figure 6.**
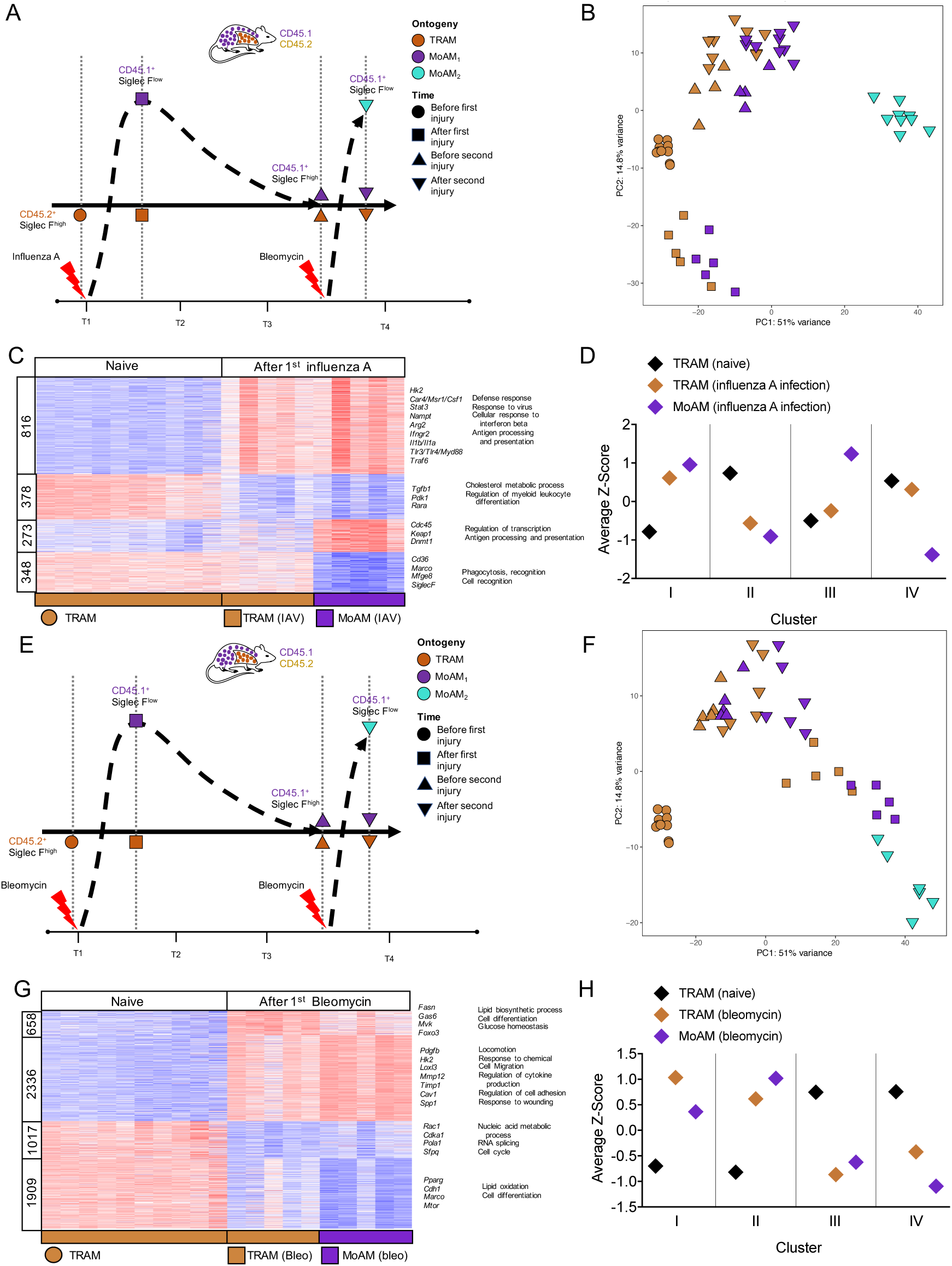
Alveolar macrophage ontogeny determines responses to influenza A virus-induced pneumonia and bleomycin-induced pulmonary fibrosis. (A) Experimental design for repeated injury experiments using influenza A virus infection as a first stimulus (see also Figure S6). (B) PCA of alveolar macrophage transcriptomes. Colors and symbols refer to panel A. (C) Heatmap shows k-means clustering of differentially expressed genes between naïve TRAM, TRAM 4 days after influenza A-induced pneumonia, and recruited MoAM at the same time (FDR < 0.01 in ANOVA-like test). Representative genes and GO processes from four clusters are shown on the right. See also Table S20. (D) Average Z-score for genes in each cluster from Panel C. Colors refer to cell populations as indicated. Z-score reflects the average gene expression within a given cluster in TRAM from naïve animals (n=10), TRAM four days after influenza A virus infection (n = 5) and MoAM recruited four days after influenza A virus infection (n=5). Mean value is shown, error bars indicate 95% confidence interval. (E) Experimental design for repeated injury experiments using intratracheal bleomycin as a first injury. Similar to A, but mice were treated with two sequential doses of intratracheal bleomycin separated by 60 days. (F) PCA of alveolar macrophage transcriptomes. Colors and symbols refer to panel E. (G) Heatmap shows k-means clustering of differentially expressed genes between naïve TRAM, TRAM 21 days after bleomycin-induced lung injury, and recruited Mo-AM at the same time (FDR < 0.01 in ANOVA like test). Representative genes and GO processes from four clusters are shown on the right. See also Table S21. (H) Average Z-score for each cluster from panel G. Colors refer to cell populations as indicated. Z-score reflects the average gene expression in a given cluster from naïve TRAM (n=10), TRAM 21 days after bleomycin (n = 5) and MoAM recruited 21 days after bleomycin (n = 5).

### Ontogeny alters the response of alveolar macrophages to influenza A virus-induced pneumonia and bleomycin-induced pulmonary fibrosis

In response to acute infection with influenza A virus (4 days) TRAM and MoAM upregulated the expression of genes involved in the immune response to virus and downregulated genes involved in macrophage homeostasis (Figure 6C, Cluster I and II). Some of these genes were exclusively upregulated or downregulated in MoAM relative to TRAM (Figure 6C, Cluster III, IV). In addition, the fold-change in the expression of almost all of these genes was more pronounced in MoAM relative to TRAM as reflected by the average Z-score for the expression of upregulated and downregulated genes in response to influenza A virus infection (Figure 6D). Similar differences in the expression of profbrotic and homeostatic macrophage genes were observed in TRAM compared with MoAM during bleomycin-induced pulmonary fibrosis (Figure 6E-H). These results highlight an important role for ontogeny in the phenotype of MoAM and TRAM in the acute phases of lung injury and fibrosis.

### The local microenvironment shapes monocyte-derived alveolar macrophages during the resolution of lung injury

We next examined the differences between MoAM and TRAM 60 days after the administration of bleomycin or influenza A infection, when lung injury is resolving. Twenty-one days after bleomycin induced lung injury, MoAM (CD45.1+) were primarily localized to areas of injury (Figure 7A). Mo-AM recruited during lung injury persisted during the resolution of influenza A or bleomycin-induced injury (60 days after administration, Figure 7B). At this time point there were significantly fewer differentially expressed genes between MoAM and TRAM irrespective of the initial injury (Figure 7C for bleomycin, Figure 7E for influenza A infection). TRAM harvested 60 days after injury had incompletely resolved their changes in gene expression induced by either influenza A infection or bleomycin administration when compared with TRAM from naïve mice. Many of the genes that differed between TRAM and MoAM during resolution still represented stimulus-specific genes and processes, a common set of differentially expressed genes was observed between MoAM and TRAM, which were similar to those we identified in normal aging. Interestingly, GO processes and genes that distinguished MoAM and TRAM at these later time points were similar irrespective of the injury (Figure 7C-F).

**Figure 7.**
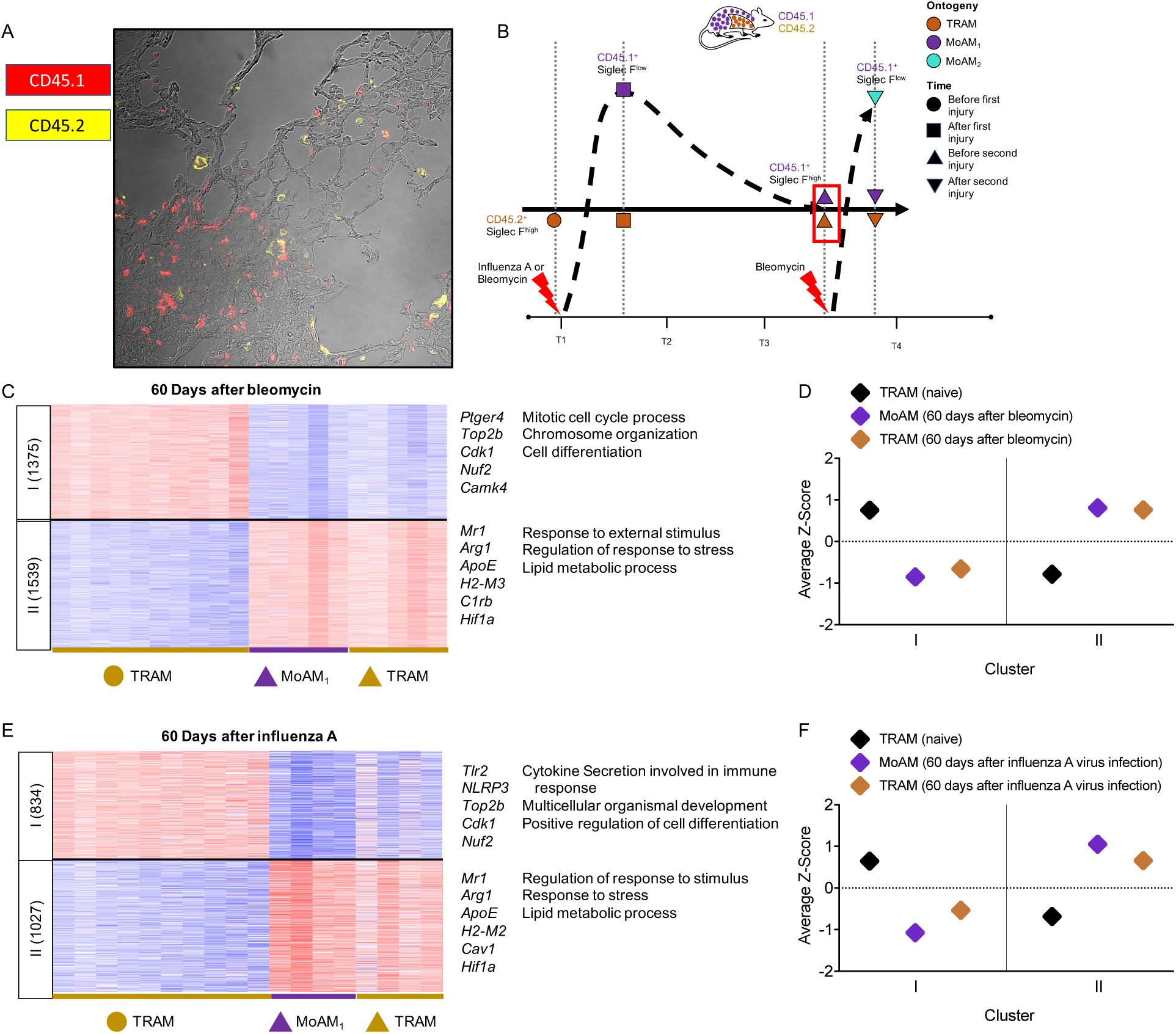
Monocyte-derived alveolar macrophages and tissue-resident alveolar macrophages become increasingly similar during the resolution of lung injury. (A) Lung sections from shielded chimeric mice 21 days after the administration of bleomycin were analyzed using immunofluorescent microscopy with antibodies against CD45.2 to label TRAM (yellow) or CD45.1 to label MoAM (red). As demonstrated in the representative section, CD45.1 MoAM were disproportionately represented in areas of injury/fibrosis compared to areas that were relatively free of fibrosis. (B) Schematic of the experimental design. The red frame indicates the populations compared in panels C-F. (C) Heatmap shows k-means clustering of differentially expressed genes (FDR < 0.01) in TRAM and MoAM retained after bleomycin-induced pulmonary fibrosis. Naïve TRAM are included as a comparison. Representative genes and GO processes are shown for each cluster (see also table S22). (D) Average Z-score for each cluster according to cell type and condition (naive TRAM, n=10, MoAM1, n=5, TRAM after bleomycin, n=5). (E) Heatmap shows k-means clustering of differentially expressed genes (FDR < 0.01) in TRAM and MoAM retained after influenza A-induced pneumonia. Naïve TRAM are included as a comparison. Representative genes and GO processes are shown for each cluster (see also table S23). (F) Average Z-Score for each cluster according to cell type and condition (naive TRAM, n=10, MoAM_1_, n=4, TRAM after bleomycin, n=4).

### The response to a second injury is independent of alveolar macrophage ontogeny

We next compared the response of TRAM and newly resident MoAM (recruited after the first injury) in response to a second dose of bleomycin (Figure 8). The response of TRAM and newly resident MoAM to bleomycin was largely similar irrespective of their ontogeny and irrespective of whether they were recruited in response to bleomycin (Figure 8A,B) or influenza A virus infection (Figure 8C,D). Despite the similarity in their responses, the differences in gene expression between TRAM and MoAM persisted during the injury. These changes were similar to the differences between TRAM and MoAM during aging and the differences between MoAM and TRAM recruited during the first injury (Figure 8E).

**Figure 8.**
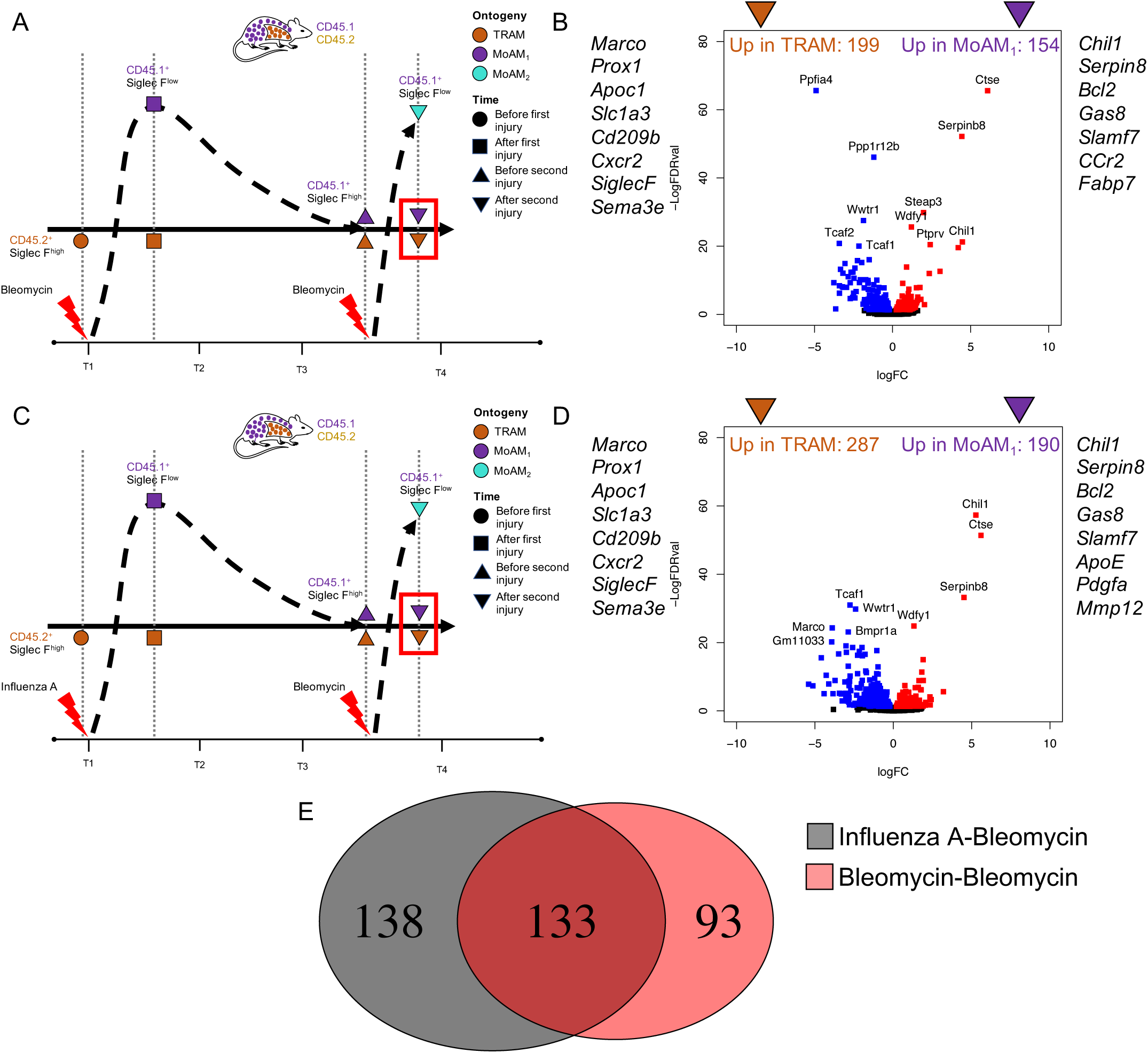
The microenvironment determines the response of alveolar macrophages to bleomycin-induced pulmonary fibrosis irrespective of ontogeny. (A) Schematic of the experimental design for panel B. (B) Volcano plot shows differentially expressed genes (FDR<0.05) between TRAM and MoAM (recruited in response to historic bleomycin exposure as the first injury) after the second injury with bleomycin. Representative genes are shown adjacent to the plot. See Table S24 for a full list of genes. (C) Schematic of the experimental design for panel D. (D) Volcano plot shows differentially expressed genes (FDR<0.05) between TRAM and MoAM (recruited in response to historic influenza A virus-induced pneumonia as the first injury) after the second injury with bleomycin. Representative genes are shown adjacent to the plot. See Table S25 for a full list of genes. (E) Venn diagram shows overlap between genes differentially expressed between TRAM and MoAM recruited in response to historic bleomycin or influenza A infection after a second injury with bleomycin. P value= 4.43e-173.

### Repeated injury to the microenvironment shapes the response of monocyte-derived alveolar macrophages during bleomycin-induced fibrosis

Repeated administration of bleomycin followed by recovery has been shown to result in more severe and persistent injury when compared with a single dose of bleomycin (Degryse et al., 2010). Consistent with these observations, we found that MoAM recruited to the lungs of animals historically exposed to bleomycin differed from those recruited to the lungs of naïve animals (Figure 9A, B). In contrast, the response of MoAM to bleomycin as a second injury was almost identical in mice historically exposed to influenza A infection and naïve mice (Figure 9C,D). A comparison of differentially expressed genes in mice historically exposed to bleomycin or influenza A virus identified genes/GO processes associated with fibrosis (Figure 9C-F). In contrast, TRAM demonstrated evidence of immune tolerance irrespective of the previous exposure (Figure 10) or ontogeny (Figure S7) as their proinflammatory response to the second injury was diminished in comparison to a naïve mouse. These results highlight the importance of both the microenvironment and ontogeny in determining the response of both MoAM and TRAM to a repeated immunologic challenge implicating the microenvironment in the development of innate immune memory.

**Figure 9.**
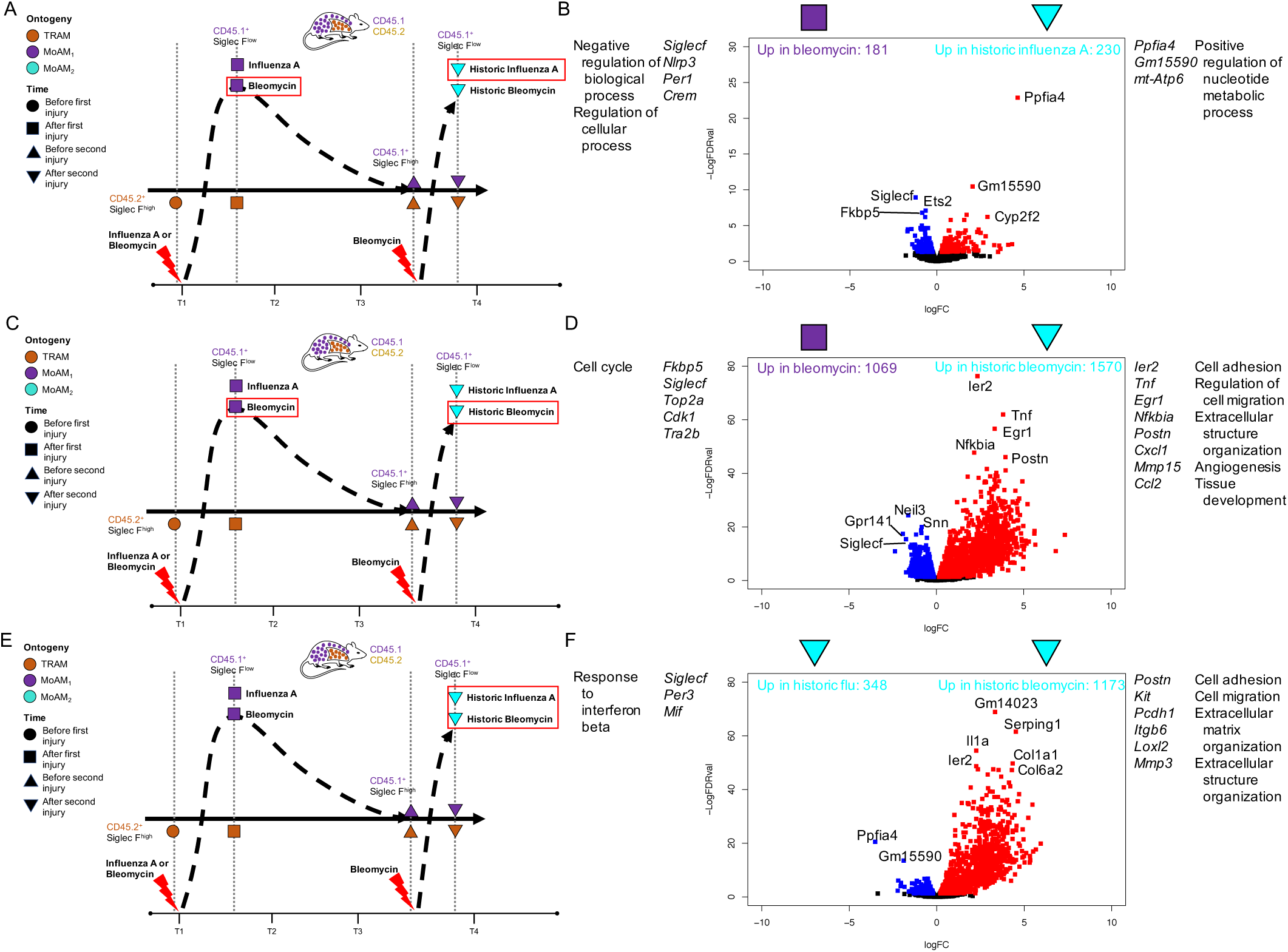
The environment determines the response of newly recruited monocyte-derived alveolar macrophages to repeated injury. (A) Schematic for experimental design for panel B. The comparison focuses on MoAM recruited to naïve lungs with those recruited to lung in historically exposed to influenza A virus. (B) Volcano plot comparing MoAM recruited to naïve lungs with those recruited to lungs historically treated with influenza A virus (FDR <0.05). Selected differentially expressed genes and GO processes are shown. See Table S26 for a full list of genes. (C) Schematic for experimental design for panel D. The comparison focuses on MoAM recruited to naïve lungs with those recruited to lungs historically exposed to bleomycin. (D) Volcano plot comparing MoAM recruited to naïve lungs with those recruited to lungs historically exposed to bleomycin (FDR <0.05). Selected differentially expressed genes and GO processes are shown. See Table S26 for a full list of genes. (E) Schematic for experimental design for panel F. Comparison between MoAM recruited in response to bleomycin in mice historically treated with bleomycin and those historically treated with influenza A infection. (F) Volcano plot comparing MoAM recruited in response to bleomycin in mice treated historically with influenza A virus or bleomycin (FDR <0.05). Selected differentially expressed genes and GO processes are shown. See Table S26 for a full list of genes.

**Figure 10.**
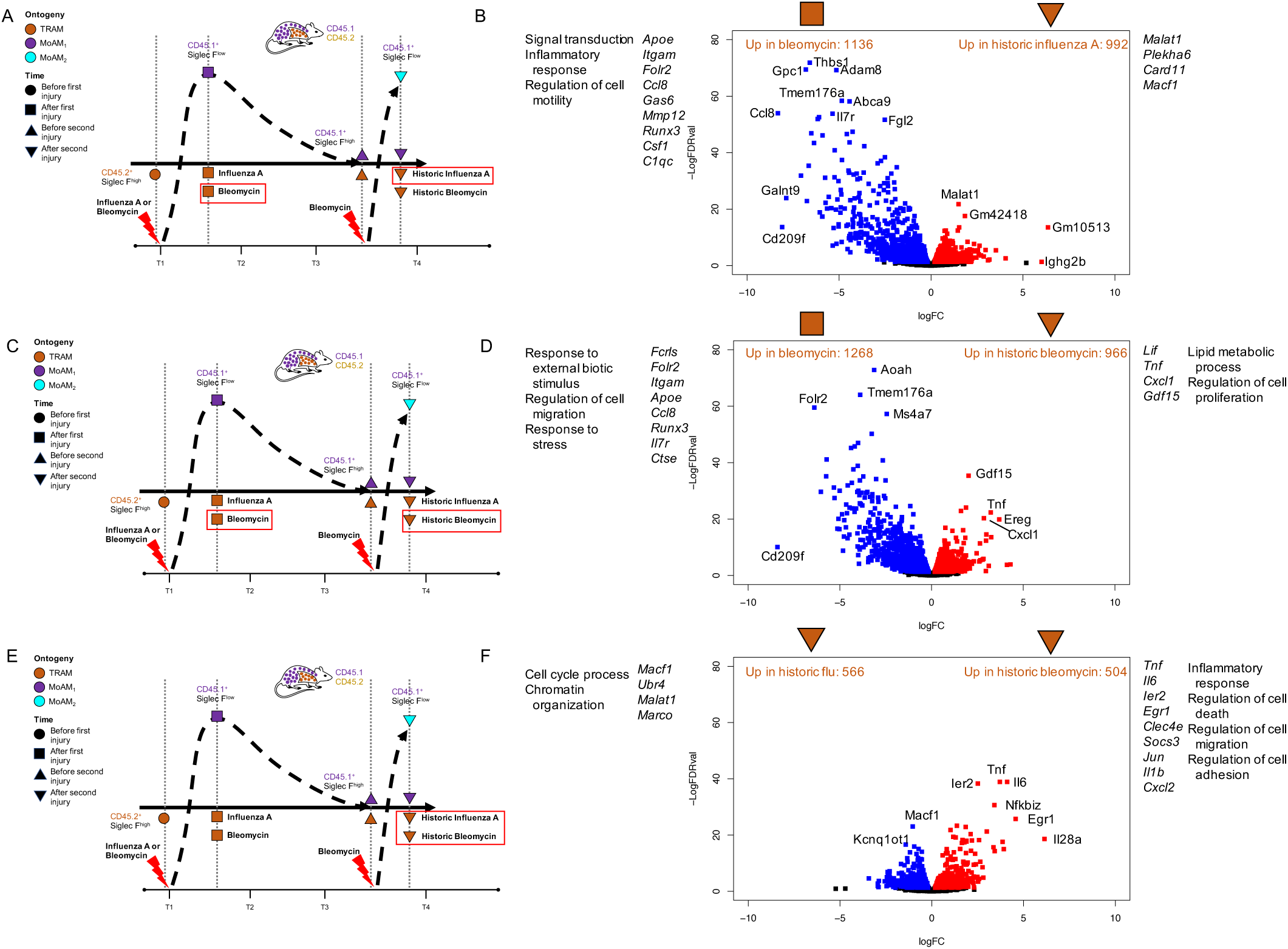
Tissue-resident alveolar macrophages demonstrate evidence of immune tolerance irrespective of the previous exposure. (A) Schematic for experimental design for panel B. The comparison focuses on TRAM from naïve lungs with those from the lungs historically treated with influenza A virus. (B) Volcano plot comparing TRAM from naïve lungs with TRAM from lungs historically treated with influenza A virus (FDR <0.05). Selected differentially expressed genes and GO processes are shown. See Table S27 for a full list of genes. (C) Schematic for experimental design for panel D. The comparison focuses on TRAM from naïve lungs with those from the lungs historically treated with bleomycin. (D) Volcano plot comparing TRAM from naïve lungs with those from the lungs historically treated with bleomycin (FDR <0.05). Selected differentially expressed genes and GO processes are shown. See Table S27 for a full list of genes. (E) Schematic for experimental design for panel F. Comparison between TRAM exposed to bleomycin or influenza A as first injury after exposure to bleomycin as a second injury. (F) Volcano plot comparing TRAM exposed to bleomycin or influenza A as first injury after exposure to bleomycin as a second injury (FDR <0.05). Selected differentially expressed genes and GO processes are shown. See Table S27 for a full list of genes.

## Discussion

Long-lived macrophage populations in the lung, brain, and liver are derived from myeloid progenitors originating from the fetal yolk sac or fetal liver and maintain their populations via self-renewal for months or years (Gomez Perdiguero et al., 2015). Depletion of tissue-resident macrophages leads to the recruitment of bone marrow-derived monocytes to the tissue where they rapidly differentiate into macrophages. Guided by signals from the local microenvironment these monocyte-derived macrophages reshape their epigenome and obtain gene expression profiles and surface markers resembling tissue-resident macrophages (Guilliams and Scott, 2017; Watanabe et al., 2019). This raises the possibility that some of the dramatic changes in the immune response to environmental challenges in aged animals are secondary to cell autonomous changes in long-lived tissue resident macrophages or retained epigenetic marks in monocyte-derived compared to tissue-resident macrophages. We used a combination of genetic lineage tracing studies and transcriptomic profiling of flow cytometry sorted alveolar macrophage populations to show that TRAM and MoAM responses to environmental challenges are instead shaped by the local microenvironment of the aged or injured lung.

Pneumonia is the most common cause of death from an infectious disease and pneumonia mortality disproportionately affects the elderly (Jain et al., 2015; Ortiz et al., 2013). Furthermore, in the year after hospital discharge, elderly survivors of pneumonia are at increased risk of developing age-related disorders including lung injury (Mittl et al., 1994), skeletal muscle dysfunction leading to immobility (Herridge et al., 2003), myocardial infarction (Corrales-Medina Vicente et al., 2012), chronic kidney disease (Murugan et al., 2010), dementia (Tate et al., 2014) and cognitive impairment (Girard et al., 2018). Therefore, interventions targeting immune dysfunction during aging are predicted to impact healthspan. We found that heterochronic adoptive transfer partially or completely reversed age-related transcriptional changes in alveolar macrophages. These findings have implications for therapy–senolytic or other novel therapies might indirectly target the immune dysfunction during aging through effects on the lung parenchyma (Baker et al., 2011; Jeon et al., 2017). Surprisingly, heterochronic parabiosis in which circulating factors and cells were shared between young and old animals, had remarkably little effect on the age-related changes in the transcriptome of either alveolar macrophages or alveolar type II cells, suggesting the alveolar microenvironment is impervious to this intervention during homeostasis.

Previous environmental challenge can prime alveolar macrophages and modulate their response to subsequent challenge, a process referred as “trained immunity” or “innate immune memory”(Netea et al., 2016). Our data suggest that innate immune memory in alveolar macrophages is conferred by the local microenvironment irrespective of ontogeny, as we detected few changes in the transcriptome or DNA methylome between TRAM and MoAM either at baseline or in response to single or repeated environmental challenges. Moreover, we found that MoAM recruited after a second exposure to bleomycin showed higher expression of pro-fibrotic genes when compared with MoAM recruited after the first injury. In sharp contrast, the response to bleomycin (second injury) was similar in MoAM and TRAM in naïve mice and mice historically infected with influenza A virus (first injury). Interestingly, TRAM exhibited immune tolerance irrespective of historic injury and ontogeny. These data are supported by physiologic data showing repeated doses of bleomycin lead to more severe and persistent fibrosis (Degryse et al., 2010). Collectively, these findings are consistent with a model in which innate immunologic memory in alveolar macrophages is driven by the lung microenvironment via non-cell autonomous mechanisms (Yao et al., 2018).

During injury TRAM and MoAM upregulated inflammatory and fibrotic genes, respectively, in response to influenza A infection or bleomycin, however, the expression of these genes was uniformly higher in MoAM. As monocyte to macrophage differentiation involves the downregulation of inflammatory genes, some of these changes are likely cell autonomous and related to macrophage differentiation (Misharin et al., 2017). Alternatively, these differences may be driven by signals from the injured microenvironment, as areas of injury were disproportionately populated by MoAM. Over time, however, TRAM and MoAM became increasingly similar. Inducible lineage tracing systems will be required to determine whether this loss of transcriptional differences between MoAM and TRAM during the resolution of injury reflects cell autonomous or microenvironment driven differentiation of MoAM (or both), or if distinct waves of monocyte-derived macrophages are recruited over the course of injury and repair (Watanabe et al., 2019).

Alveolar macrophages avidly take up inhaled environmental particles small enough to access the alveolar space, which are then transported to the mouth via mucociliary clearance and excreted in the feces (Semmler et al., 2004). Interestingly, we found that myeloid cells are not recruited in response to short term relatively high level exposure to inhaled urban particulates. These data are consistent with observations in patients after lung transplantation. Even though transplant rejection and enhanced susceptibility to infection in these patients likely creates an environment more inflammatory than normal, alveolar macrophages in the transplanted lung are almost exclusively derived from the donor, even years after transplant (Nayak et al., 2016). Interestingly, we have shown that both influenza A infection and the intratracheal administration of bleomycin cause a depletion of TRAM (Misharin et al., 2017). Based on these findings, and the observed replacement of TRAM by MoAM after radiation or the intratracheal administration of liposomal clodronate, it is tempting to speculate that the retention of MoAM in the lung requires depletion of the TRAM pool and opening of the alveolar niche (Guilliams and Scott, 2017). If so, this would explain our observation that adoptive transfer of TRAM was impossible without prior depletion with liposomal clodronate.

Our study has some limitations and important negative findings. First, while the reproducible changes in the alveolar macrophage transcriptome with aging are attributable to the local microenvironment of the alveolus, the mechanisms by which those interactions change with aging are not known, and are likely multiple. Our data and growing single-cell RNA-Seq atlases should allow the identification of putative receptor ligand pairs associated with macrophage differentiation in aging alveolar macrophages or alveolar type II cells that might be targeted to “rejuvenate” old macrophages. Second, ontogeny may play a role in shaping the proteome independent of the transcriptome and since we measured only a small number of surface proteins via flow cytometry we would be unlikely to detect these changes. In addition, we observed reproducible transcriptomic differences between MoAM and TRAM that did not affect the response to bleomycin or to influenza A infection. We tested whether this regulation involves changes in DNA methylation using reduced representation bisulfate sequencing. These findings were negative as we did not find any association between the transcriptional changes and differences in the DNA methylome, nor could we detect differences in DNA methylation within putative enhancer regions reported in alveolar macrophages. It is therefore likely that the transcriptional differences between TRAM and MoAM are maintained via different epigenetic mechanisms, such as histone modifications, as has been previously described (Gosselin et al., 2014; Gosselin et al., 2017; Lavin et al., 2014).

In conclusion, our study highlights the overwhelming importance of the lung microenvironment in shaping the transcriptional response of alveolar macrophages during aging and in response to repetitive injury. During acute injury when alveolar macrophage differentiation is incomplete and the microenvironment is abnormal, cellular ontogeny is key to the function of alveolar macrophages, but these differences are transient. Studies with parabiosis suggest the age-related changes in the alveolar microenvironment occur independently of circulating factors or immune cells. These findings dramatically expand the possible mechanisms by which immune function might change during aging and suggest that therapies that directly target the aging microenvironment, for example metformin, senolytics or others, might improve macrophage function via non-cell autonomous mechanisms.

## METHODS

### CONTACT FOR REAGENT AND RESOURCE SHARING

Further information and requests for resources and reagents should be directed to and will be fulfilled by the Lead Contact: Scott Budinger.

### EXPERIMENTAL MODEL AND SUBJECT DETAILS

#### Mice

All animal experiments and procedures were performed according to protocols approved by the Institutional Animal Care and Use Committee at Northwestern University. C57BL/6J (Jax 000664) and CD45.1 (Jax 002014) mice were bred in our facility and our colonies are refreshed yearly with mice purchased from the Jackson Laboratory. When indicated, young and aged C57BL/6 mice from NIA NIH colony were used. Number of animals per group was determined based on our previous publications. Investigators were not blinded to the group allocation. Mice were housed at the Center for Comparative Medicine at Northwestern University, in microisolator cages, with standard 12 hr light/darkness cycle, ambient temperature 23C and were provided standard rodent diet (Envigo/Teklad LM-485) and water ad libitum.

#### Murine Model of PM Exposure

Inhalational exposure to PM2.5 CAPs was performed as previously described (Chiarella et al., 2014). Briefly, mice were housed 8 hr per day for 3 consecutive days in a chamber connected to a Versatile Aerosol Concentration and Exposure System (VACES). We exposed control mice to filtered air in an identical chamber connected to the VACES in which a Teflon filter was placed on the inlet valve to remove all particles. We estimated ambient PM_2.5_ concentrations as the mean of reported values from the 4 EPA monitoring locations closest to our location. The mean concentration in the PM exposure chamber was 118.3 ± 5.21 mg/m^3^.

#### Adoptive transfer of alveolar macrophages

Mouse primary alveolar macrophages were isolated by bronchoalveolar lavage performed on euthanized mice with 3 ml of PBS with 1 mM EDTA. Only male mice were used as a source of alveolar macrophages. The lavage was centrifuged at 300 g for 10 min and resuspended in RPMI supplemented with 10% FBS and plated in a density of 100,000 cells/cm^2^. Alveolar macrophage purity was analyzed by flow cytometry and was confirmed to be >95%. Recipient mice were pretreated with 50 ul of clodronate-loaded liposomes (Liposoma) 72 hours before the adoptive transfer of donor alveolar macrophages to partially deplete tissue-resident alveolar macrophages and make the niche permissive for engraftment. Donor alveolar macrophages (1.5×10^5^ cells in 50 µl of PBS) were transferred via intratracheal instillation into isoflurane anesthetized mice. CD45.1/.2 mice were used to discriminate between the recipient and donor cells.

#### Bone marrow chimeras and bone marrow chimeras with thoracic shielding

Bone marrow chimeras were established by transferring 5 × 10^6^ bone marrow cells isolated from C57BL/6 mice (this strain expresses CD45.2 alloantigen) into 8-wk-old lethally irradiated (single dose of 1,000 cGy γ-radiation using a Cs-137–based Gammacell-40 irradiator; Nordion) recipient mice (expressing CD45.1 alloantigen). Mice were maintained on autoclaved water supplemented with antibiotics (trimethoprim/sulfamethoxazole; Hi-Tech Pharmacal) for 4 wk after bone marrow transfer and then switched to normal housing regimen. CD45.2 to CD45.1 bone marrow chimeras were used for experiments 8 wk after bone marrow transfer. 8 wk after bone marrow transfer, >95% of all leukocytes and 100% of monocytes and neutrophils in peripheral blood were of donor origin. Bone marrow chimeras with thoracic shielding were used to assess the origin of pulmonary macrophages (TRAMs vs. Mo-AMs) and were generated in a manner similar to that previously described (Janssen et al., 2011; Janssen et al., 2010), with additional modifications (Misharin et al., 2014; Misharin et al., 2017). Briefly, to protect TR-AMs from radiation, we applied a uniform lead shield that covered the lungs during irradiation. To eliminate the residual recipient bone marrow in the shielded region mice were treated with myeloablative agent busulfan (30 mg/kg body weight; Sigma-Aldrich)(Chevaleyre et al., 2013) 6 h after the irradiation, followed 12 h later by bone marrow infusion. Chimerism was assessed 2 mo after the procedure via FACS analysis of the peripheral blood collected from facial vein. Bone marrow chimeras with thoracic shielding were maintained on antibiotics for 4 week as described above and then switched back to the normal housing regimen.

#### Parabiosis

Parabiosis was performed as described previously (Kamran et al., 2013). Briefly, parabiont mice were co-housed for 2 weeks prior to surgery. After anesthesia and preparation of the surgery site, the skin flap from elbow to knee was created on left and right sides of parabionts, accordingly, ankles and knees of parabionts were joined using non-absorbable suture and skin flaps were joined using continuous uninterrupted suture. Mice were treated with Buprenorphine SR during the first two weeks of recovery and provided with recovery diet gel (ClearH_2_O). After recovery mice were switched back to the normal housing regimen.

#### Tissue preparation and flow cytometry

Tissue preparation for flow cytometry analysis and cell sorting was performed as previously described (Misharin et al., 2013; Misharin et al., 2017). Blood was collected into EDTA-containing tubes via facial vein bleed (from live animals) or cardiac puncture (from euthanized animals). Whole blood was stained with fluorochrome-conjugated antibodies, and erythrocytes were then lysed using BD FACS lysing solution. For single-cell suspension obtained from tissues erythrocytes were lysed using BD Pharm Lyse, and cells were counted using Nexcelom K2 Cellometer C automated cell counter (Nexcelom) with AO/PI reagent. Cells were stained with eFluor 506 (eBioscience) viability dyes, incubated with FcBlock (BD), and stained with fluorochrome-conjugated antibodies (see STAR Methods). Data and cell sorting were performed at Northwestern University RLHCCC Flow Cytometry core facility on SORP FACSAria III, BD LSR II, BD Fortessa and BD Symphony instruments (Becton Dickinson). Sorting was performed using a 100-µm nozzle and 40 psi pressure. Compensation, analysis and visualization of the flow cytometry data were performed using FlowJo software (Tree Star). “Fluorescence minus one” controls were used when necessary to set up gates.

#### Transcriptome profiling via RNA-Seq

Flow cytometry sorting was used to isolate mouse alveolar macrophages and AT2 cells at indicated time points for each experiment. Cells were sorted with MACS buffer, pelleted and lysed in RLT Plus buffer supplemented with 2-mercaptoethanol (Qiagen). The RNeasy Plus Mini Kit (Qiagen) was used to isolate RNA and remove genomic DNA. RNA quality was assessed with the 4200 TapeStation System (Agilent). Samples with an RNA integrity number (RIN) lower than 7 were discarded. RNA-seq libraries were prepared from 100ng of total RNA using the NEB Next RNA Ultra Kit (Qiagen) with poly(A) enrichment. Libraries were quantified and assessed using the Qubit Fluorimeter (Invitrogen) and 4200 TapeStation. Libraries were sequenced on NextSeq 500 instrument (Illumina) at 75 bp length, single end reads. The average reading depth across all experiments exceeded 6×10^6^ per sample and over 94% of reads had a Q-score greater than 30. For RNA-seq analysis, reads were demultiplexed using bcl2fastq (version 2.17.1.14). Read quality was assessed with FastQC. Samples that did not pass half of the 12 assessed QC statistics were eliminated. Low quality basecalls were trimmed using trimmomatic (version 0.33). Reads were then aligned to the *Mus musculus* reference genome (with mm10 assembly) using the TopHat2 aligner (version 2.1.0). Counts tables were generated using HTSeq (version 0.6.1). Raw counts were processed in R (version 3.4.4) using edgeR (version 3.20.9) to generate normalized counts. Negative binomial likelihood with default setting followed by generalized linear models fitting was used to estimate differentially expressed genes. FDR q-values were used to correct for multiple comparisons and a value of 0.05 was used as a threshold to indicate statistical significance. K-means clustering was performed using the built-in R stats package (version 3.4.4). Gene Ontology analysis was performed using GOrilla (Eden et al. 2009) on two unranked gene lists. The RNA-seq datasets are available at GEO: GSE134397. Computationally intensive work was performed on Northwestern University’s Quest High-Performance Computing Cluster (Northwestern IT and Research computing).

#### Modified reduced representation bisulfite sequencing

Measurement and analysis of DNA methylation was performed as previously described (McGrath-Morrow et al., 2018; Singer, 2019; Walter et al., 2018; Wang et al., 2018; Weinberg et al., 2019). Briefly, genomic DNA was isolated from sorted macrophage populations (Qiagen AllPrep Micro kit). Endonuclease digestion, size selection, bisulfite conversion, and library preparation were prepared as previously described from approximately 250 ng of input genomic DNA. Bisulfite conversion efficiency was 99.6% as estimated by the measured percentage of unmethylated CpGs in λ-bacteriophage DNA (New England BioLabs N3013S) spiked in to each sample. Equimolar ratios of four libraries per run were sequenced on NextSeq 500 sequencer (Illumina), at 75 bp length, single end reads using NextSeq 500/550 V2 High Output reagent kit. Demultiplexing, trimming, alignment to *Mus musculus* reference genome (with mm10 assembly), and methylation calling were performed as previously described. Quantification was performed using the DSS R/Bioconductor package (Feng et al., 2014) and the SeqMonk platform (Andrews, 2018). The RRBS dataset is available at GEO: GSE134238. Computationally intensive work was performed on Northwestern University’s Quest High-Performance Computing Cluster (Northwestern IT and Research computing).

#### Statistics

Data were reported as mean±SEM and subjected to 1-way ANOVA. For pairwise significance, we applied two tailed Student’s t-test with Benjamini-Hochberg correction for multiple comparisons (R: stats: version 3.4.4; Prism 7: graphpad). Statistical methods for RNA-seq were described above. Sample power and size were estimated based on our previous work and published papers. The observed data met or exceeded required sample size criteria. The statistical parameters and criteria for significance can be found in figure legends.

#### Data and software availability

The RNA-seq datasets, containing raw and processed data, are available at GEO: GSE134238 (RRBS dataset), GSE134397 (RNA-seq dataset). R code that was used for the RNA-seq and RRBS analysis is available on GitHub.

## ACKNOWLEDGMENTS

A.V.M.: NHLBI HL135124, NIH NIAMS AR061593, ATS/Scleroderma Foundation Research Grant, DOD grant PR141319, and BD Bioscience Immunology Research Grant. P.A.R.: HL076139. A.B.: NIH HL125940 and the Thoracic Surgery Foundation, Society of University Surgeons, John H. Gibbon Jr. Research Scholarship from American Association of Thoracic Surgery. B.D.S.: HL128867, Parker B. Francis Research Opportunity Award. J.I.S.: NIH AG049665, HL048129, HL071643, HL085534. G.M.M.: NIH ES015024 and ES025644 and ES026718. K.R.: NIH HL079190 and HL124664. G.R.S.B.: NIH ES013995, HL071643, AG049665, VA BX000201, DOD PR141319. ES015024 H.P.: NIH AR064546, AG049665 HL134375 and the Mabel Greene Myers Chair. N.S.C.: NIH grants AG049665, HL071643, CA197532. Core Facilities (Flow Cytometry and Genomics NCI Cancer Center Support Grant P30 CA060553), the Feinberg School of Medicine, the Center for Genetic Medicine, and Feinberg’s Department of Biochemistry and Molecular Genetics, the Office of the Provost, the Office for Research, and Northwestern Information Technology, and maintained and developed by Feinberg IT and Research Computing Group. S.W.: MSD Life Science Foundation, Public Interest Incorporated Foundation, Japan. David W. Cugell and Christina Enroth-Cugell Fellowship Program H.P.: NIH grants AR064546, HL134375, AG049665, and UH2AR067687, the United States-Israel Binational Science Foundation (2013247), the Rheumatology Research Foundation (Agmt 05/06/14), Mabel Greene Myers Professor of Medicine and generous donations to the Rheumatology Precision Medicine Fund. C.J.G: NIH grant HL143800. Z.R. Driskill graduate program in life science. A.B NIH HL125940, HL145478, HL147290, HL147575 K.M.R. NIH HO071643, HL128194,GM096971 W.E.B. NIH AG049665 R.M. NIH AG049665

The authors declare no competing financial interests.

## AUTHOR CONTRIBUTIONS

Conceptualization, G.R.S.B., A.V.M.; Methodology, G.R.S.B., A.V.M., Z.R., and A.C.M.P.; Investigation, A.C.M.P., Z.R., N.J., S.W., M.C., Z.L., L.S., J.M.D, R.P., S.S., C.E.R., K.A.H and A.Y.M.; Resources, M.A., H.A.V., A.B., S.M.B., M.J., and A.G.; Data Curation, Z.R.,T.S., P.A.R, K.R.A, K.N. and B.D.S; Writing – Review & Editing, G.R.S.B., A.V.M., B.D.S, E.R.W, C.J.G, K.M.R., N.S.C, J.I.S, W.E.B., G.M.M, A.B., H.P., R.I.M.,Z.R.; Visualization, Z.R.,G.R.S.B., and A.V.M.; Supervision, G.R.S.B., A.V.M.; Project Administration and Funding, G.R.S.B. and A.V.M.

---

**Figure S1.**
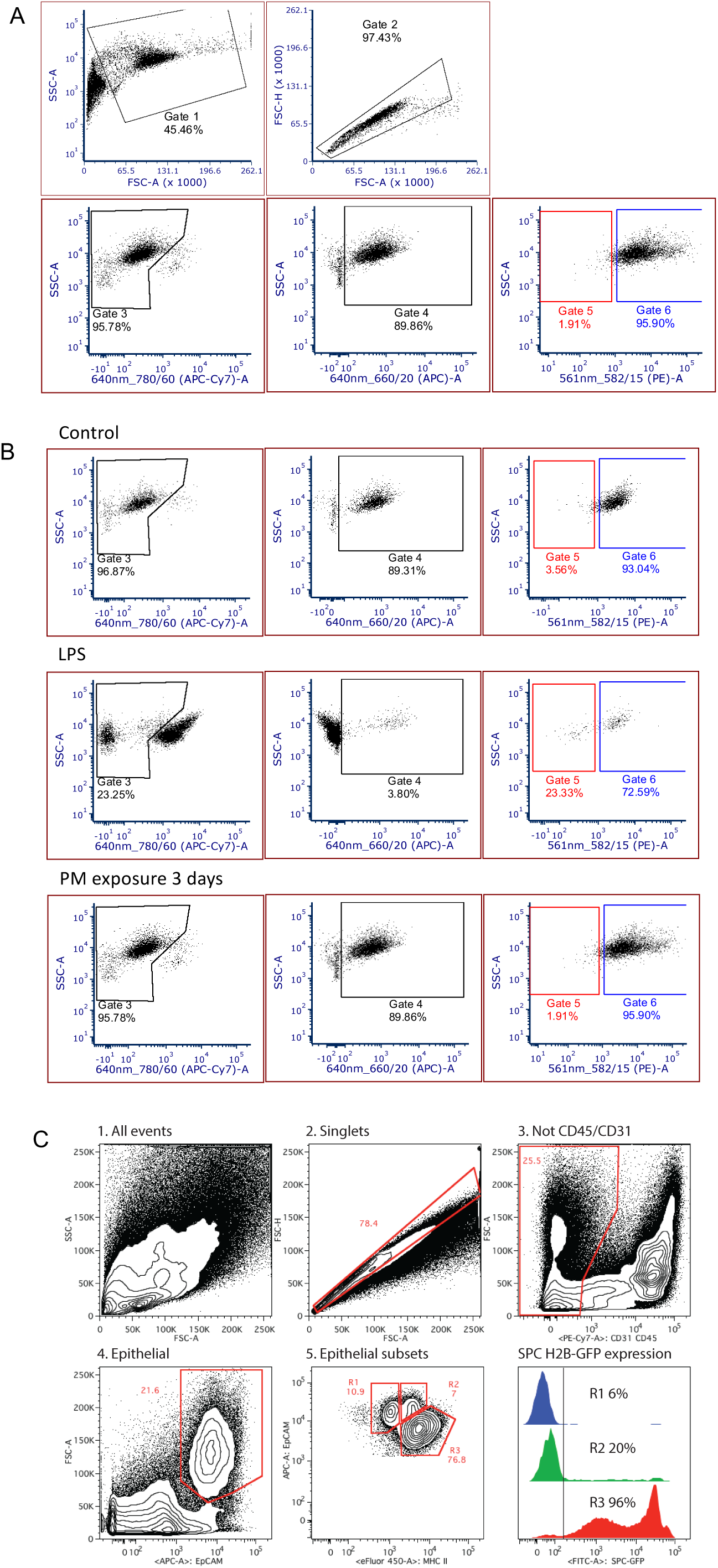
Gating strategy for labeling alveolar macrophages with PKH26 and isolation of alveolar type II cells via flow cytometry. (A) Alveolar macrophages were labeled by intratracheal administration of PKH26 *in vivo*. After 24 hours, animals were treated intratracheally with PBS (50 µL), LPS (1 mg/kg in 50 µL PBS) or were exposed to concentrated particulate matter air pollution (6 hours of exposure ∼100-120 µg/m^3^, 6 hours daily for 3 consecutive days). Concentrations of particles are estimates based on particle measures from the inlet and outlet of the concentrator and reported ambient PM_2.5_ measures at nearby monitoring stations. Alveolar macrophages were collected via bronchoalveolar lavage 72 hours after the first exposure and stained with the following reagents for flow cytometry: APC/Cy7 anti-mouse Ly-6G, Alexa Fluor 647 anti-mouse F4/80 and Helix NP Green live/dead. (B) Representative flow plots for a single animal in each experimental group. (C) Validation of gating strategy for isolation of alveolar type II cells from mice. Single cell suspensions were prepared from the lungs of BAC transgenic mice expressing GFP driven by the Surfactant C (SFTPC) promoter (*Tg-SFTPC-H2B-GFP*) mice and analyzed via flow cytometry. Epithelial cells were identified as singlets/live/CD45-negative/CD31-negative/EpCAM-positive cells and further subdivided based on EpCAM and MHC II expression. R1: EpCAM-high MHC II-negative, R2: EpCAM-high MHC II-positive and R3: EpCAM-intermediate MHC II-positive. Over 95% of cells in gate R3 were positive for surfactant protein C-GFP reporter and this gate was used to identify alveolar type II cells during conventional flow sorting.

**Figure S2.**
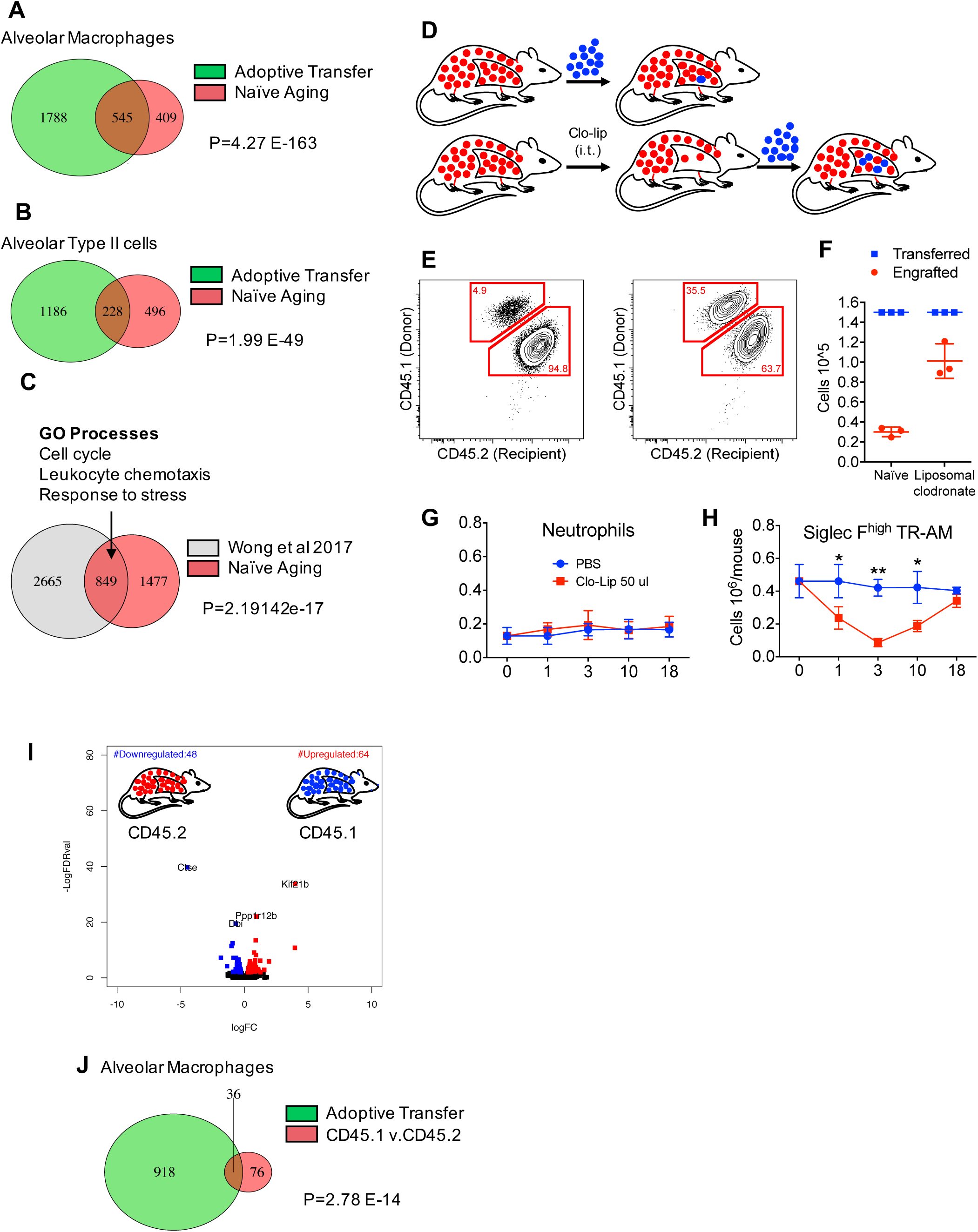
Heterochronic adoptive transfer of TRAM using CD45.1/CD45.2 mice. (A) Venn diagram shows overlap between genes differentially expressed in TRAM from naïve young (4 months of age) and old (18 months of age) mice and in adoptive transfer experiments (Refers to Figure 2A and Figure 2F). (B) Venn diagram shows overlap between genes differentially expressed in alveolar type II cells from naïve young and old mice and in adoptive transfer experiments (Refers to Figure 2A and Figure 2H). (C) Venn diagram shows overlap between differentially expressed genes between young and old TRAM in this study (Fig. 2A) and those reported by Wong et. al. (D) Schematic design of heterochronic adoptive transfer experiments with or without intratracheal liposomal clodronate. (E) Representative flow cytometry plots from CD45.2 mice treated without (left panel) or with (right panel) liposomal clodronate (25 µL, intratracheally) before the adoptive transfer of 1,5×10^5^ TRAM from CD45.1 mice. (F) Quantification of adoptively transferred TRAM 60 days after adoptive transfer (n=3 mice per group). (G) Mice were treated with liposomal clodronate 25 µL intratracheally and the number of neutrophils was measured in the single cell suspension prepared from the lung using flow cytometry. (H) Mice were treated with intratracheal liposomal clodronate 25 µL intratracheally and the number TRAM (SiglecF^high^) was measured using flow cytometry. (I) TRAM were isolated from the lungs of CD45.1 (Jax 002014) or CD45.2 mice (Jax 000664) via flow sorting, RNA was isolated and analyzed using RNA-Seq. Differentially expressed genes (FDR < 0.01) are shown (see also Table S3 for full list of genes). (G) Venn diagram shows overlap between differentially expressed genes in CD45.1/CD45.2 TRAM and genes differentially expressed in heterochronic adoptive transfer experiments (Refers to Figure 2F).

**Figure S3.**
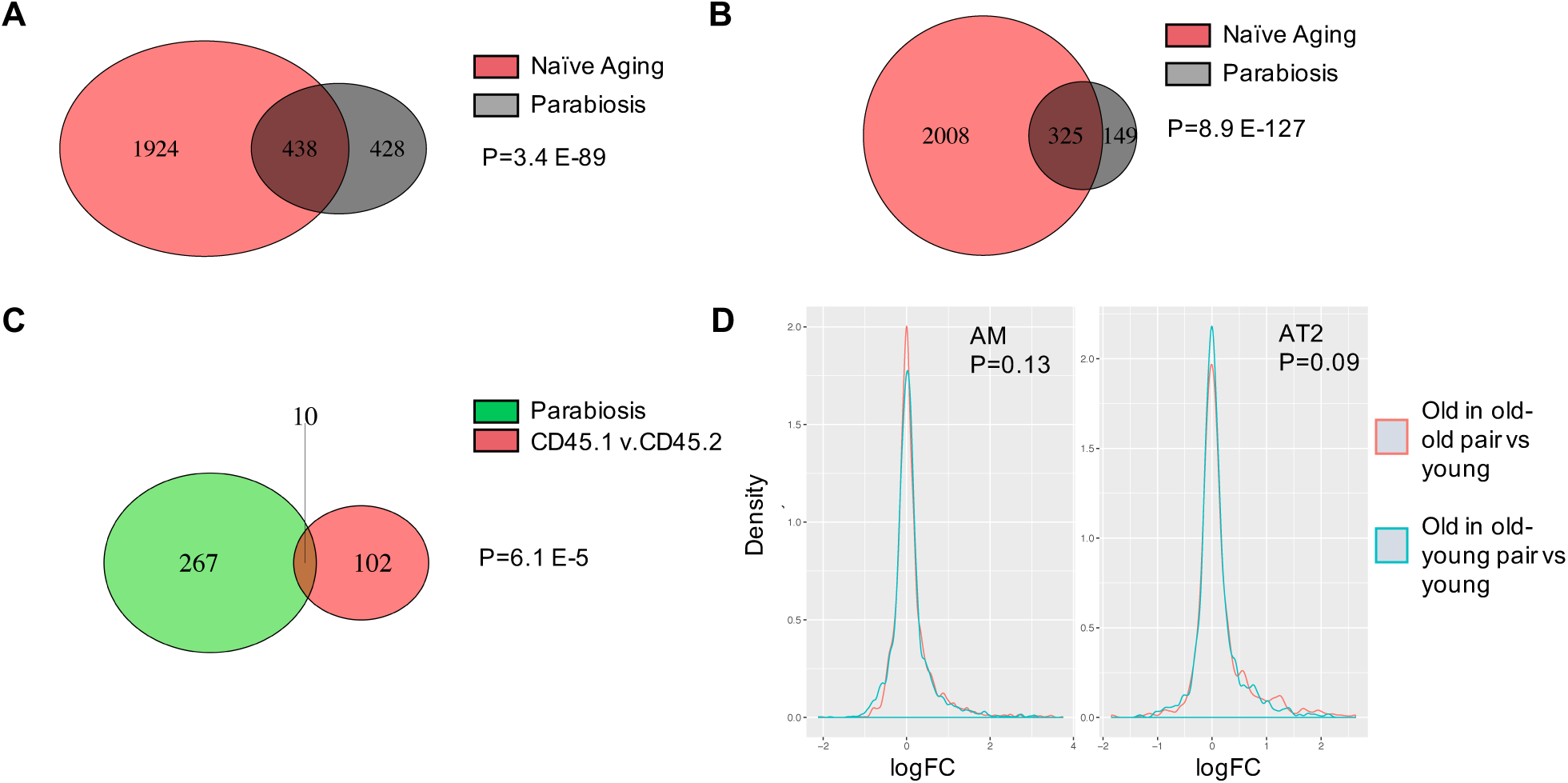
Parabiosis does not reverse age-related transcriptional changes in the lung. (A) Venn diagram showing overlap of differentially expressed genes in young/young compared with old/old parabiont pairs and differentially expressed genes in TRAM identified between 4 month and 18 month old mice aged in our colony (from Figure 2). (B) Venn diagram showing overlap of differentially expressed genes in young/young compared with old/old parabiont pairs and differentially expressed genes in alveolar type II cells identified between 4 month and 18 month old mice aged in our colony (from Figure 2). (C) Venn diagram showing overlap of differentially expressed genes in TRAM from parabionts in young/young compared with old/old pairs with differentially expressed genes (with FDR <0.01) in TRAM from CD45.1 and CD45.2 alveolar macrophages. (D) Distribution of log-fold change showing no significant difference (p>0.05 by Wilcoxon rank sum test) for differentially expressed genes in TRAM and AT2 identified between 6 month and 18 month-old mice aged in our colony (from Figure 2) and heterochronic parabiont pairs.

**Figure S5.**
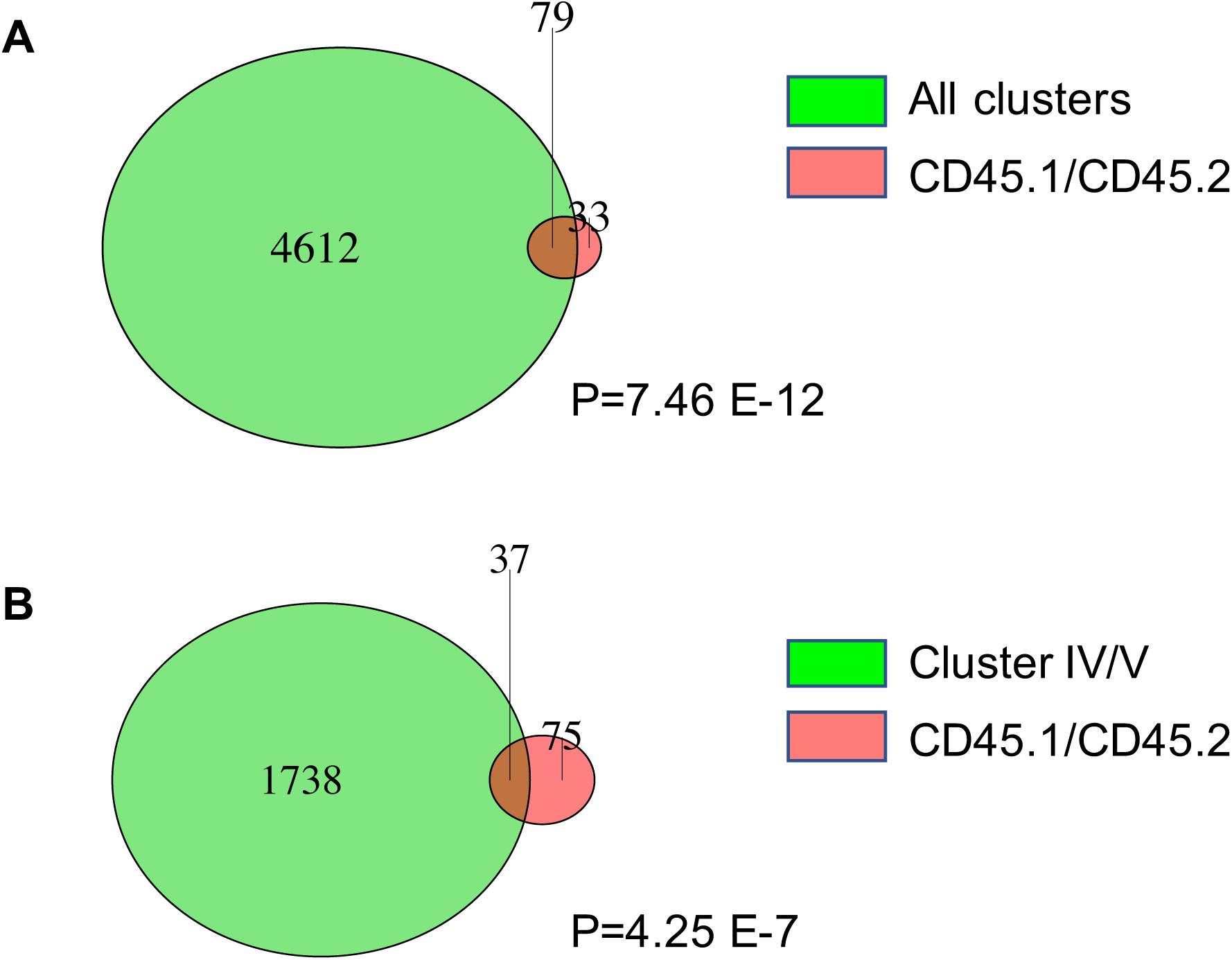
Differences in CD45.1/CD45.2 strains do not explain differential gene expression in tissue-resident alveolar macrophages and monocyte-derived alveolar macrophages from aged shielded chimeric mice. (A) Venn diagram showing overlap between all differentially expressed genes in TRAM and MoAM from shielded chimeras in aging (See Figure 5A) with those differentially expressed in TRAM from untreated CD45.1 and CD45.2 mice. (B) Venn diagram showing overlap between differentially expressed genes in Clusters IV and V of Figure 5A (comparison of TRAM and MoAM in aging shielded chimeras) with those differentially expressed in TRAM from untreated CD45.1 and CD45.2 mice.

**Figure S6.**
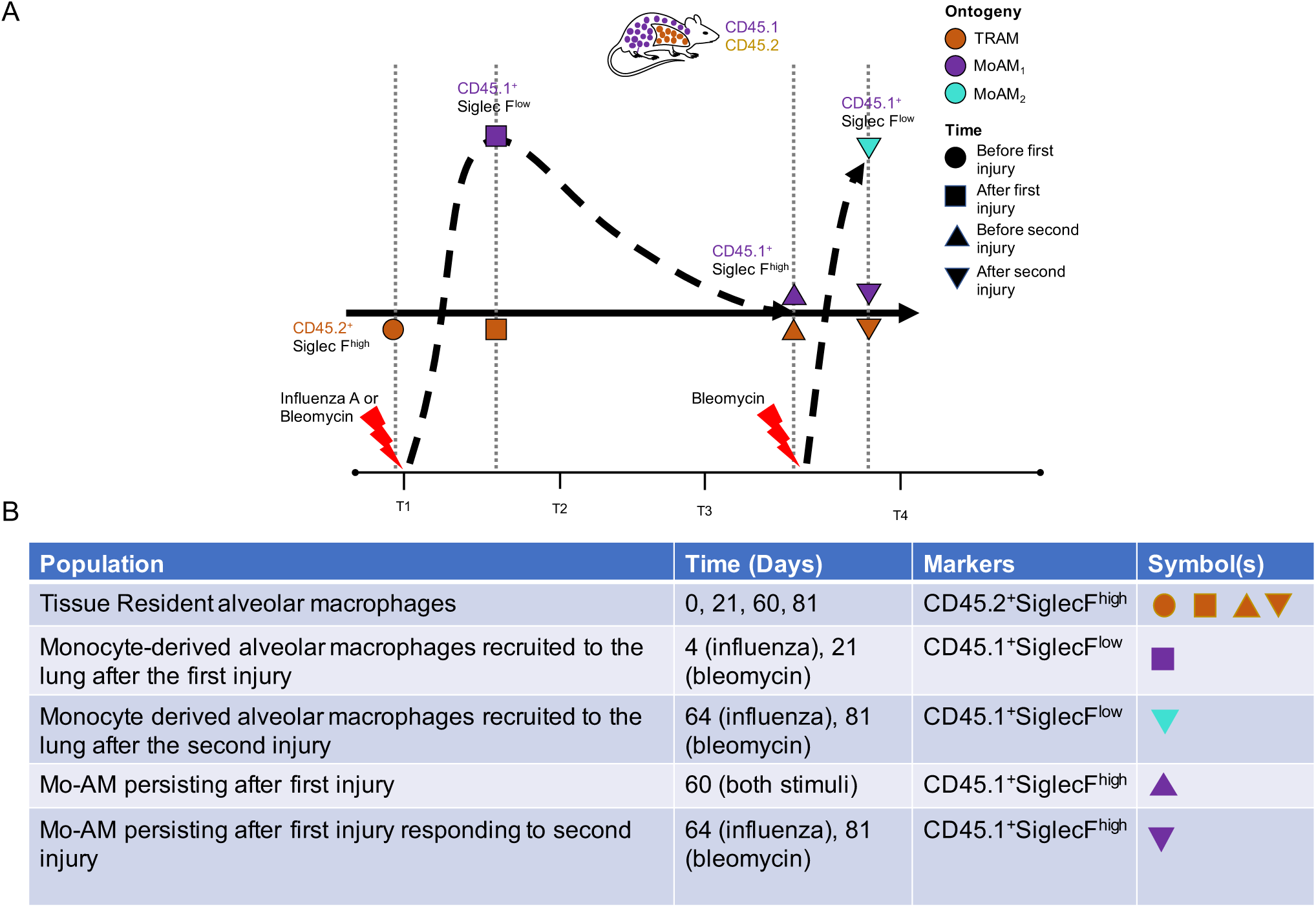
Strategy to examine differential responses in MoAM and TRAM. (A) Experimental design. Mice were administered either intratracheal bleomycin or infected with influenza A virus on day 0 followed by treatment with bleomycin on day 60. Alveolar macrophage populations were collected as indicated. Refers to Figure 6. Flow cytometry gating was performed as previously described (see Misharin et al., 2017). Table describes flow markers used to sort cell populations for RNA-Seq in Figure 6.

**Figure S8.**
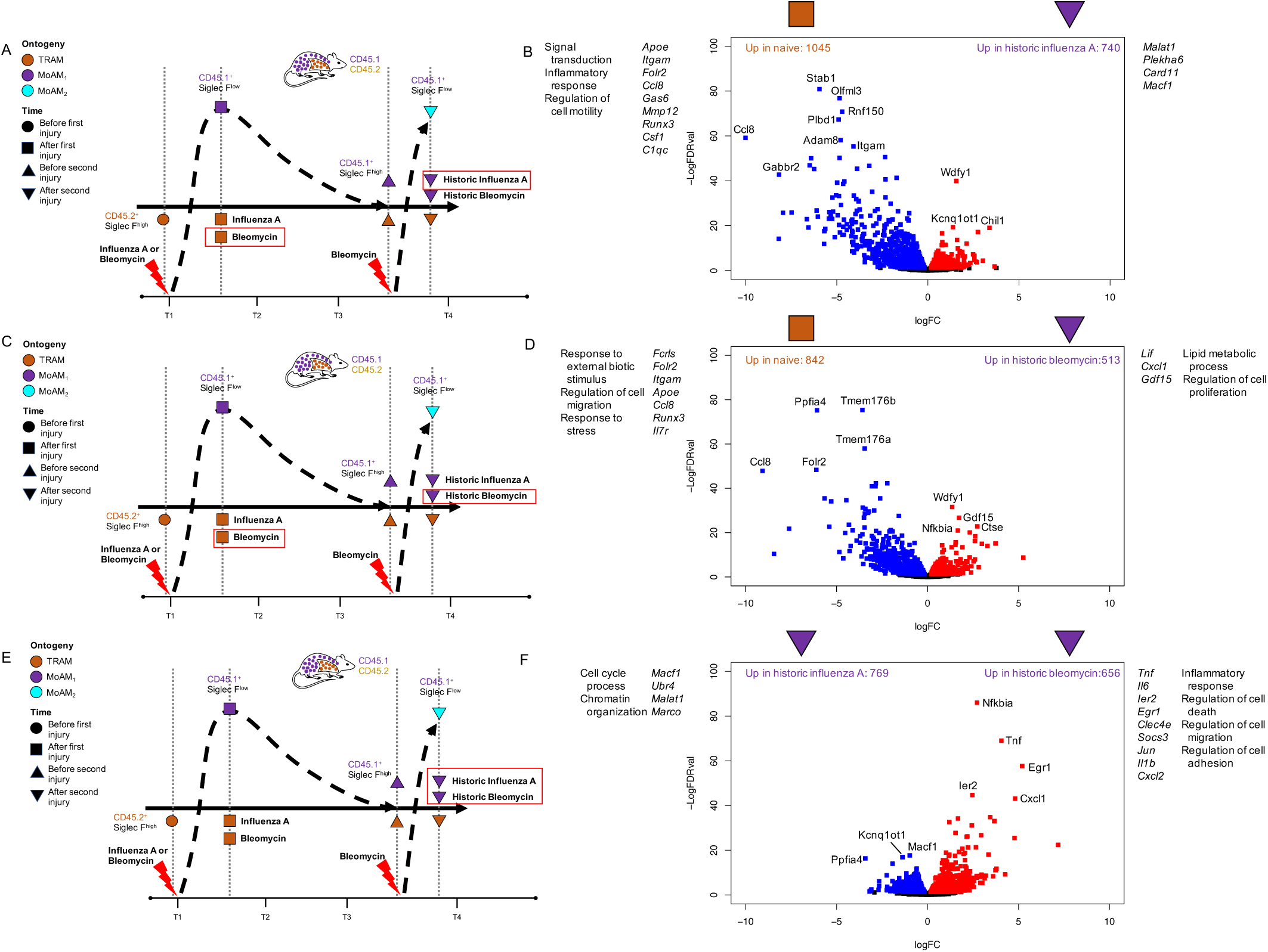
Tissue-resident alveolar macrophages demonstrated evidence of immune tolerance irrespective of ontogeny. (A) Schematic for experimental design for panel B. The comparison focuses on TRAM during the first injury (bleomycin) and MoAM recruited after historic exposure to influenza A virus and re-challenged with bleomycin. (B) Volcano plot comparing TRAM during the first injury (bleomycin) and MoAM recruited after historic exposure to influenza A virus and re-challenged with bleomycin (FDR <0.05). Selected differentially expressed genes and GO processes are shown. See Table S28 for a full list of genes. (C) Schematic for experimental design for panel D. The comparison focuses on TRAM during the first injury (bleomycin) and MoAM recruited after historic exposure to bleomycin and re-challenged with bleomycin. (D) Volcano plot comparing TRAM during the first injury (bleomycin) and MoAM recruited after historic exposure to bleomycin and re-challenged with bleomycin (FDR <0.05). Selected differentially expressed genes and GO processes are shown. See Table S28 for a full list of genes. (E) Schematic for experimental design for panel F. The comparison focuses on MoAM recruited after the historic exposure to bleomycin or influenza A virus and re-challenged with bleomycin. (F) Volcano plot comparing MoAM recruited after the historic exposure to bleomycin or influenza A virus and re-challenged with bleomycin (FDR <0.05). Selected differentially expressed genes and GO processes are shown. See Table S28 for a full list of genes.

